# A chemical genetic screen uncovers novel seed priming agents capable of persistent perturbation of anthocyanin regulation in *Arabidopsis thaliana*

**DOI:** 10.1101/2023.05.22.541227

**Authors:** Katrina M. Hiiback, Malcolm M. Campbell

## Abstract

Priming is a general term for a phenomenon in which exposure to an early environmental stimulus results in a more rapid or vigorous response when the plant is exposed to subsequent challenges. Various types of priming have been described: ‘systemic acquired resistance’ against biotic stimuli, ‘epigenetic stress memory’ induced by repeated abiotic stimuli, and ‘seed priming’ treatments which apply water and additional adjuncts to seeds to improve crop performance. Using a high-throughput chemical genomic approach, thousands of small molecules were screened to identify compounds capable of ‘chemical priming’, asking if these could serve as artificial environments to persistently induce altered response to later abiotic challenges when applied as seed treatments. Attenuation of expected visual anthocyanin accumulation was chosen as a screening phenotype due to the visual nature of these pigments, as well as their biological roles in development and stress response. Several novel structural categories of molecules were identified that had the ability to reduce total anthocyanin accumulation in 7-18-day old seedlings induced by later low temperature challenge, persisting days after the removal of the compounds. Application variables were explored with thought to future use of the priming compounds as functional treatments: a dose-dependent relationship was established, additional effects on growth and development were documented to ensure minimal detrimental side effects on the treated plants, and the necessary temporal window of treatment was explored and reduced. Cross testing of the priming treatments identified by low temperature screening for ability to reduce anthocyanin induction by alternative exogenous and endogenous conditions showed consistent attenuative effect and revealed that the effect of priming on this metabolic phenotype was not specific to low temperature response. The presented research represents a proof-of-concept for the functional potential of seed priming with novel compounds, and highlights anthocyanin accumulation as a flexible component of plant stress response.

## Introduction

Plants are sessile organisms, and as such must develop internal mechanisms for responding to external environmental conditions throughout their lives from germination to senescence. The sum of the interactions between the expressed genotype of an individual and their external environment produces an overall phenotype, which may then affect growth and survival [1–3]. An extensive body of literature exists examining responses to individual acute abiotic environmental stimuli, with a growing number of studies addressing combinations of stress stimuli both biotic and abiotic, but the mechanisms which govern responses to multiple or repeated challenges can be difficult to predict and are still not fully understood [4–6].

Plant priming as a general concept describes the processes in which a plant is exposed to an initial environmental stress stimulus, recovers, then displays a faster, more vigorous or otherwise altered response to additional challenges thereafter [7–10]. These phenomena have been of great research interest, as understanding and development of these priming mechanisms may represent an opportunity to improve plant performance under adverse conditions with minimal cost to growth and fitness [11–13]. Different mechanisms have been reported as contributing to primed responses, dependent on context. Against biotic stimuli, accumulation of dormant pattern-activated protein kinases, chromatin modifications, and messengers like azelaic acid have been described [14,15]; against abiotic stimuli, epigenetic mechanisms are frequently implicated [16,17] and physical stalling of transcriptional machinery at relevant genes has been observed [18]. Hormonal signaling and metabolic regulation have been shown to contribute to primed phenotypes as well [19–23]. The duration of altered primed responses is also variable, dependent on context: for examples, a one-time ABA treatment applied to *Vicia faba* seedlings induced salinity resistance up to 8 days later [24]; *Arabidopsis* thaliana seedlings exposed to different regimes of either constant or pulsed low temperature treatment resulted in alteration of cold acclimation patterns and transcriptional response also over a timespan of days [25]; drought exposure during the seedling stage, stem elongation stage or both in *Triticum aestivum* resulted in longer-lasting results improving tolerance to further drought and heat during grain filling weeks later [26]. Many priming strategies are therefore possible when considering the possible types of initial stimulus or stimuli to be used, the timing of their introduction to the plant, and the output phenotype or measure of interest.

Seed priming is a technique with a long history, with the conceptual basis having been described in some form as early as the mid-19th century [27], performed to increase germination efficiency and improve seedling vigour. In its most basic form, seeds are soaked (‘hydropriming’); additional strategies may be used to control the rate of water update and seed hydration (e.g. solid matrix priming, ‘osmopriming’ in osmotic solutions); with further common additions to the priming solution generating further corresponding technological terms (e.g. plant hormones – ‘hormopriming’, mineral nutrients – ‘nutripriming’, microbial inoculation – ‘biopriming’) [28,29]. ‘Chemical priming’ has been used as a catch all term for studies applying additional natural and synthetic compounds with known properties to seeds, including the fungicide and synthetic hormone paclobutrazol; molecules with functions related to signalling and stress including β-aminobutyric acid (BABA), melatonin, the nitric oxide donor sodium nitroprusside (SNP), and reactive oxygen species hydrogen peroxide (H_2_O_2_); various antioxidants, nanoparticles and more [30–33]. In addition to the observed germination benefits, it has been demonstrated that the effects of seed priming can extend beyond germination to grant plants greater resilience against later abiotic stress conditions, as recently observed against drought conditions at two vegetative growth phases in wheat [34].

Seed priming and other plant priming phenomena therefore share an implication of some form of plant ‘memory’, in which information from an early experience is retained by the plant and alters response to subsequent environmental experiences [10,35]. Although many molecular agents have been used in different chemical priming techniques applied to seeds and to adult plants, with few exceptions the existing literature focuses on the use of compounds which are either endogenous to plants, already known to have bioactivity such as synthetic hormones or fungicides, and/or are complex mixtures such as the total seaweed extracts used in biostimulant assays [30,36–38]. Therefore, the full breadth of possible molecular space capable of provoking persistent response has yet to be explored.

Plant chemical genetics is a growing field of study based on the concept of using large numbers of low-molecular weight compounds to perturb and probe biological pathways, similar to pharmacological drug discovery [39,40]. Small molecules which can alter phenotypes of interest may uniquely provide insight into multifaceted environmental response pathways, either by acting as general antagonists which can overcome systemic redundancy, or as specific agonists, stimulating a particular element such as to highlight its role in the overall response [41]. Significant successes have been achieved applying chemical genomic strategies to identify active compounds affecting hormone signalling and endomembrane trafficking, even if the precise mode of action has not yet been elucidated [42]. Targeted screening experiments have also been applied to identify agents capable of metabolic inhibition, reducing lignification [43] or to mimic a stress response, phosphate starvation [44]. A chemical genomic screen can thus be understood as a range of novel chemical micro-conditions that can be applied to uncover or create phenotypic responses in ways both broad and specific that mutations in forward genetic screens may not, leading to greater understanding of the system and species of interest.

Anthocyanin accumulation was chosen as the primary output phenotype for a chemical genomic assay seeking compounds capable of altering abiotic stress responsiveness in *Arabidopsis thaliana*. Anthocyanins are pigmented phenylpropanoid secondary metabolites which are ubiquitous across most taxa of land plants, can be produced by all types of plant tissues and combined with their related upstream flavonoids make up a family of over 9000 molecules [45,46]. Their biosynthetic pathway has been well characterized and is coordinately activated by upregulation of enzyme gene transcription by a MYB-bHLH-WDR (MBW) regulatory complex [47,48]. The production and accumulation of anthocyanins and flavonoids is triggered by many developmental, hormonal, metabolic and environmental factors including sugar signalling and abiotic stress responses [49,50,59,51–58] and regulated with a high degree of specificity by a large network of repressors [60,61]. This endogenous fine-tuning capacity enables and emphasizes the importance of the functional roles of these metabolites throughout the lifespan of a plant. The accumulation of anthocyanin pigments induced by *A. thaliana* seedling low temperature response was therefore ideal to leverage as a visual output to facilitate high-throughput screening of a combinatorial compound library.

This research thus assessed if a chemical genetic approach could be applied to the concept of chemical priming: to identify novel molecular agents capable of altering later seedling phenotype when applied solely during the process of seed imbibition and germination, ideally continuing to affect phenotype not only after treatment removal but persisting through an additional low-temperature challenge. After identification of several candidate structural categories of priming treatments capable of altering anthocyanin accumulation, highly similar structural analogues were obtained for structure function comparison and dose-response analyses. The necessary duration of priming treatment exposure and possible light interaction were explored with thought towards potential future utility as practical treatments. Additional growth parameters were assessed in primed plants transplanted to soil and grown until senescence to provide evidence that the priming dose capable of persistently altering low-temperature-induced anthocyanin induction in seedlings did not also impose a significant cost or trade-off on normal *A. thaliana* development. The flexibility of treatments to attenuate anthocyanins in different inductive contexts was explored, and found that the chemical priming treatments were more likely to be persistent anthocyanin regulators than responsive to a specific abiotic stress. As the molecular mechanisms underlying all forms of priming are still not yet comprehensively understood, identification and optimization of treatments capable of serving as highly specific ‘chemical environments’ to consistently trigger persistent phenotypic and metabolic responses will be useful tools to enable further study.

## Materials and Methods

### Plant materials, growth conditions and low temperature challenge

*Arabidopsis thaliana* Col-0 seeds were surface sterilized in 10% bleach and 1% Triton X-100 for 25 minutes, then rinsed five times with equal volumes of sterile deionized water. 8-10 sterilized seeds and 200 µl liquid MS growth media were distributed into each well of 96-well microplates. Standard liquid MS media used throughout consisted of 1X Murashige and Skoog basal medium ([62]; Caisson Labs), 0.05% MES buffer (2-((N-morpholino) ethanesulfonic acid), 1% sucrose, and 0.1% Gamborg vitamin solution ([63]; Sigma) in Milli-Q purified water adjusted to pH 5.8 as used in similar chemical screening studies [64,65]. Further experimental treatments were added directly to the liquid growth media as described below.

Plant materials were grown in several different Conviron growth cabinets at 21°C, with 16-hour light / 8-hour dark long-day photoperiod and ∼133-176 µmol m^-2^ s^-1^ light intensity. Low temperature exposure for compound screening was performed in a Percival growth chamber on the same long-day photoperiod schedule held at -1°C with ∼80-100 µmol m^-2^ s^-1^ light.

### Chemical screening: library composition and chemical analogue selection

The chemical library used for screening was a custom diversity-oriented library of 4182 compounds designed by members of the Centre for the Analysis of Genome Evolution and Function (CAGEF) at the University of Toronto, selected from the NOVACore and EXPRESS-Pick collections provided by the ChemBridge corporation (San Diego, CA). Purchased 10 mM stock solutions in dimethylsulfoxide (DMSO) were diluted to 2.5 mM working stocks with additional DMSO (Bioshop) and stored in foil-sealed 96-well conical-bottom plates at -20°C until use.

Prepared wells of seeds in liquid media in a 96 well plate were each treated as a unique chemical microenvironment by individual addition of 2 µL screening compound per well for a final treatment concentration of 25 µM (or 2 µL of DMSO as control) similarly to protocols from previous studies [64,65]. Plates were sealed with micropore tape and stored at 4°C for 3-4 days for stratification, then moved to lighted growth conditions. After 3 days of growth, treatment media was removed, seedlings were rinsed by adding and removing a 200 µl aliquot of standard MS media, then replenished with a second 200 µl aliquot of standard MS media. At 8 days of growth, plants were moved to low temperature treatment previously observed to reliably generate a visible anthocyanin accumulation phenotype (-1°C for 10 days with 1% sucrose supplementation; Supplementary Figure 1). Plant tissue was photographed in-plate using a Canon EOS Rebel XT and photographs were initially qualitatively examined for individual wells showing phenotypic alteration (reduced purple colouration) vs. controls.

Additional structural analogues of hit compounds identified in the low temperature assay were identified and obtained based on structural similarity and catalog availability using the Hit2Lead tool (Chembridge, http://www.hit2lead.com/).

### Anthocyanin extraction and measurement

Plant tissue for anthocyanin extraction was collected by pooling 2-3 wells containing 8-10 seedlings each into samples of fresh weight 10-50 mg. n ≥ 3 pooled samples were collected for all data. Samples were dried carefully to remove residual liquid media, then stored at -80°C in 1.5 mL microtubes. Frozen tissue was ground within the storage microtubes using sterilized microtube pestles under liquid nitrogen. Anthocyanin extraction and quantification were performed as described by Neff and Chory [66] with modifications to the scale of the protocol to account for small sample mass. Acidified methanol (1% HCl) was added to tissue and incubated at 4°C overnight; to remove chlorophyll, an equal volume of distilled water and 2.5x volume chloroform were added, mixed, then centrifuged at 15000 rpm 2-5 minutes for phase separation; the upper aqueous phase was analyzed with a NanoDrop 1000 or 2000 spectrophotometer measuring absorbance at 535 (A_535_) and 657 nm (A_657_). A_535_ – A_657_ was reported as final absorbance (A) per gram (g) of starting fresh weight (A/g).

### Dose-response assay for priming compound confirmation

Seeds were grown in liquid MS media in 96 well plates with chemical seed priming treatment and subsequent low temperature treatment as described previously, with modification of the concentration of priming treatment to assess the relationship between treatment dose and phenotypic response. Preparations of the identified small molecule treatments were diluted from stock solutions such that application of 2 µL of each screening compound in DMSO provided final treatment concentration in growth media of 5 µM, 10 µM, 25 µM, 50 µM, and 100 µM, expanding from and replicating the original screening concentration of 25 µM. 2 µL DMSO was applied as control. Seedlings were collected at 8 days growth prior to chilling and after the subsequent 10 days of low temperature treatment. This dose-gradient experimental design was performed in triplicate.

### Soil-grown lifespan morphometric analysis

Seeds were stratified and grown in liquid MS media in 24-well microplates (all volumes doubled) with DMSO or 25 µM chemical treatment in DMSO applied for 7 days then removed by rinsing as described previously. At 10 days old, seedlings were transferred to soil for further growth observation. Individual seedlings from each treatment condition at the same growth stage (first set of true leaves approximately 1mm in diameter) were planted in each quadrant of square 4” nursery pots filled with Sunshine Mix #1 soil (Sun Gro Horticulture, Agawam MA). Pots were grown in 16-hour light, 8-hour dark long-day conditions at 21°C with light intensity of approximately 110-120 µmol m^-2^ s^-1^ . Half of the pots of each seed treatment group (n = 6-8 individuals per priming treatment) were subjected to a low temperature treatment at 20 days of growth: flats were moved from 21°C to 4°C (photoperiod maintained, light intensity approximately 72-116 µmol m^-2^ s^-1^) for a period of 3 days, then returned to 21°C for the remainder of the experimental period. The remaining pots (n = 6-8 per priming treatment) were maintained under control growth conditions. Each priming treatment was assessed in triplicate experiments.

Observations of floral transition (visible emergence of an inflorescence), rosette diameter, primary inflorescence length and dry biomass were guided by the morphometric analysis presented by Boyes *et al.* [67]; quantitative measures were performed manually with a standard ruler marked in millimeters. For above-ground dry biomass, plants were allowed to fully senesce before cutting below the rosette for sampling; samples were subsequently dried overnight at 60°C before mass measurements were taken. For observation of S1 germination, the same liquid media microplate set-up was performed with no additional chemical treatment and 4 days cold stratification prior to moving to light; seedlings were observed for germination (radicle emergence) one day afterwards.

### Reduction of chemical treatment duration – modifications to set-up and analysis

Seeds were grown in liquid MS media in 96-well plates with 25 µM chemical seed priming treatment and subsequent low temperature treatment as described previously, with several modifications to the duration and timing of seed treatment exposure. In the first, priming treatment was applied as usual during plate set-up, remained on seeds throughout 3 days 4°C stratification, and rinsed with two aliquots of standard MS media upon moving the plates to lighted 21°C growth conditions. In the second, plates were set up without priming treatment during stratification; priming treatments were applied when plates were moved to lighted growth conditions, remained on seeds for 3 days during germination, then rinsed with two aliquots standard MS at day 3 of growth as performed previously.

A subsequent modified priming timetable was designed to reduce chemical seed priming duration to a single day. Each day during the 3-day light-grown germination seedlings were harvested and assessed. Plates were again set up without priming treatment during stratification; priming treatments were applied either immediately when plates were moved to lighted growth conditions, after 1 additional day, or after 2 additional days. In all cases, treatment remained on seeds for one day; seedlings were rinsed the day following application with two aliquots of standard MS growth media as performed previously. Each modified priming regime experiment was performed in triplicate

### Low nitrogen chemical priming assay

Experiments examining the effect of chemical priming of seedlings subjected to subsequent nitrogen deprivation was performed as described in Naik (2016) [68] with minor modifications. In brief, seeds were surface sterilized and 8-10 seeds were sown into 200 µL sterile MS media with 0.05% MES buffer, 0.1% Gamborg vitamin solution and 0.33% sucrose in 96 well plates with either 2 µL chemical priming treatment in DMSO or 2 µL pure DMSO as control for final treatment concentration of 25 µM. Prepared plates were wrapped in foil and stratified at 4°C in the dark for 4 days to ensure uniform germination, then moved to long-day conditions (16 hours light / 8 hours dark) at 21°C for 3 days. Priming treatments were removed by pipette and seeds were rinsed with an aliquot of untreated fresh MS media. At this stage, seedlings received either a new 200 µL aliquot of the previously described MS media (nitrogen-sufficient (N+) control condition) or 200 µL nitrogen-deficient (N-) media. The nitrogen deficient condition medium consisted of modified N-free ½ strength MS [62], 0.05% MES buffer, 10 mM sucrose, 10 mM ammonium nitrate (NH4NO3) and 0.1% Gamborg vitamin solution [63] adjusted to pH 5.7, representing a 5:1 C:N ratio. Seedlings were grown for an additional 7 days prior to anthocyanin extraction and observation.

### *pap1D* overexpression line chemical priming assay

pap1D homozygous T-DNA insertion line seeds overexpressing *PAP1 (PRODUCTION OF ANTHOCYANIN PIGMENT 1)* were obtained as a gift from the Rothstein lab at the University of Guelph. This line was originally generated and described by Borevitz *et al.* [69], obtained from the ABRC (accession CS3884) and confirmed by true phenotypic presentation over two generations. pap1D seeds were sterilized as described previously for wild-type Col-0 and 8-10 seeds per well were sown into liquid MS media in 96 well plates. Chemical priming compounds were added to the liquid media as previously described at a concentration of 25 µM and remained on seeds throughout 4 days stratification and 3 days germination, removed by rinsing then replenishment with fresh untreated MS media. Anthocyanin observations were made 7 days after the removal of compounds without additional abiotic treatment.

### High exogenous sucrose chemical priming assay

Standard liquid MS media consisting of 1X Murashige and Skoog basal medium ([62]; Caisson Labs), 0.05% MES buffer, and 0.1% Gamborg vitamin solution ([63]; Sigma) in Milli-Q purified water adjusted to pH 5.7 was amended for this assay. 30, 100 or 200 mM sucrose was added to the standard MS liquid preparation for the elevated sucrose assay (where 30 mM (1%) was the previously established baseline for this experimental protocol). These amended media were added to wells with 8 day old wild type Col-0 seedlings previously primed as described for screening either with DMSO or 25 µM BAO treatment by removing the prior media by pipette, rinsing the seedlings with new media, then adding a full fresh 200 µL aliquot of the given concentration to each test well. Tissue collection for anthocyanin extraction and observation was performed every three days at late afternoon thereafter.

### Statistical analysis

All data presented were analyzed using GraphPad Prism 9 (Dotmatics, San Diego, USA). Two-way ANOVA was performed to compare each chemical priming treatment with the DMSO control within each stress treatment (before/after low temperature; N+/N-; at each observation time point for other data). P <0.05 is reported as significant (*), with additional annotation for p < 0.01 (**), <0.001(***), <0.0001(****) as indicated on figures (Dunnett’s multiple comparison test). EC50 approximations over incomplete dose-response curves were performed with non-transformed treatment concentrations using GraphPad Prism 9 default equation “[Inhibitor] vs response” (Hill’s standard slope =1, asymmetrical profile-likelihood ratios for confidence intervals), with assumption that A/g value for the DMSO control (dose = 0) in each grouping were the maximum with a value of zero entered as hypothetical minimum. For 1-day compound priming data, two-way ANOVA was performed with Tukey’s multiple comparison test to distinguish significance within and between variables (timing of priming treatment x control treatment x low temperature exposure).

## Results and Discussion

### Chemical screening: testing secondary abiotic stimuli, establishing low-temperature anthocyanin accumulation phenotype and initial qualitative compound identification

Several abiotic stress stimuli (temperature extremes, pH extremes, high light intensity, high nitrogen (N)) were initially evaluated for their effect on *A. thaliana* seedling phenotype when grown in a liquid MS well plate system, designed to enable future chemical treatment addition. Different degrees of each stressor were applied to optimize environmental treatment for the growth system, with the ideal of identifying a non-lethal visible phenotype to facilitate high-throughput screening which emerged gradually over time so that persistence of the chemical priming treatment could be evaluated. Most abiotic treatments evaluated (pH variations, high temperature) resulted in seedlings displaying a chlorosis-like phenotype over a period of one-two days, except for low temperature and low nitrogen (Naik, 2016), which both resulted in anthocyanin accumulation. Exposure to low temperature (-1°C – 1°C) in media supplemented with 30 mM sucrose (1%) induced seedlings to produce visible accumulation of red-purple anthocyanin pigmentation after 8-10 days (Fig S1), consistent with previous evidence linking low temperature and anthocyanin production in the presence of sucrose [49,53]. Due to the gradual duration of anthocyanin induction, the strong visual aspect of the phenotype for high throughput evaluation, and potential overlap in involvement with response to other abiotic challenges such as high light and nutrient availability [70], this regime and target phenotype were selected for chemical library screening.

A custom library of 4182 synthetic small molecules optimized for bioavailability [71] was screened to tentatively identify small molecules which, when applied to seeds throughout stratification and germination and subsequently removed, were able to persistently alter the expected phenotype typically induced by later low temperature treatment (Fig 1). Screening compounds were applied at 25 µM for seven days total treatment. Many chemical priming treatments were identified in an initial round of screening which perturbed post-chilling phenotype, with the largest number (61 compounds) producing seedlings with a consistently green attenuated-anthocyanin phenotype. This reduction of the expected visible red-purple anthocyanin accumulation after the abiotic challenge was initially assumed to be a proxy for resilience or resistance to low temperature – i.e. plants which did not mount this protective response did not require it. The 61 compounds from preliminary screening were re-evaluated with an identical qualitative round of secondary screening, after which 8 compounds again produced a green phenotype (Fig S2). Of these 8 compounds, five possessed a chemical structure in which an oxime group was attached to a phenyl ring (four to benzene forming a benzaldehyde oxime (BAO) moiety, one to naphthalene forming a naphthaldehyde oxime (NAO) moiety) suggesting the importance of this substructure. Interestingly, searching the full library initially screened showed that four additional BAO compounds were included in the primary screen and were not identified by their effect on anthocyanin phenotype. To further investigate the efficacy of the identified oximes and begin to establish structure function relationships in the context of this chemical priming phenotype, additional analogues as available were obtained for both BAO and NAO base structures, and several other hit compounds (HC) were carried forward, including one thiazole (TZ) (Table 1).

**Figure 1:**
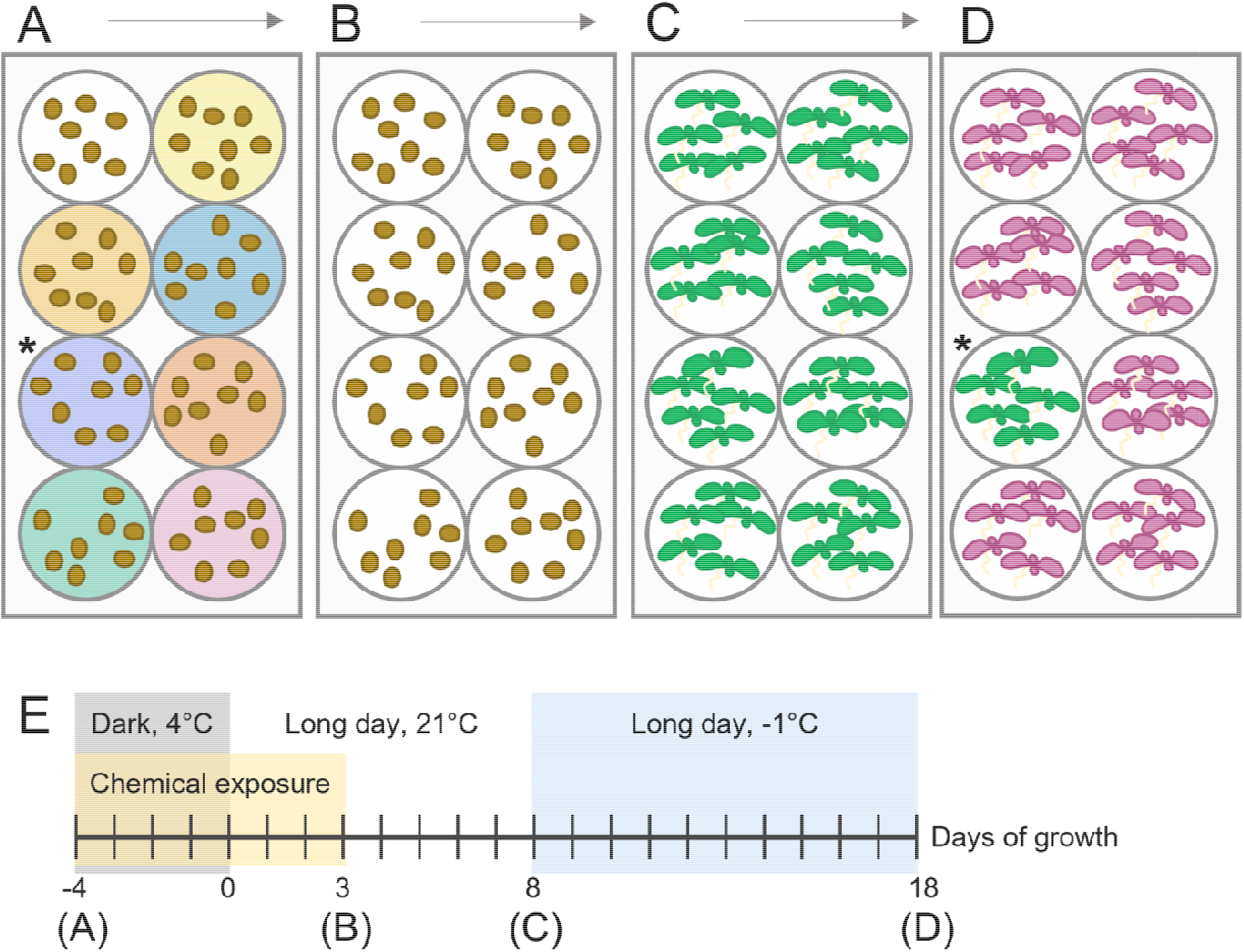
Chemical seed priming process used for library screening and subsequent analyses. (A) 8-10 *A. thaliana* Col-0 seeds were surface sterilized, then sown in well plates containing liquid MS growth media supplemented with sucrose and either priming treatments in DMSO solution or DMSO alone as control. (B) After 4 days of stratification (24h dark, 4°C) and 3 days growth in long day conditions (16h light / 8h dark, 21°C), the priming treatment media was removed, the seedlings rinsed, then wells replenished with a fresh aliquot of standard growth media. (C) At 8 days growth, the cotyledons and first true leaves have emerged for most seedlings, and the plates are moved from 21°C to -1°C. (D) After 10 days at -1°C, anthocyanin pigments are observable in wild-type control seedlings. At this time, wells in which the chemical priming treatment altered environment response were visible due to anthocyanin attenuation and their resulting green phenotype (*). (E) shows timeline with steps A-D indicated in summary.

**Table 1:**
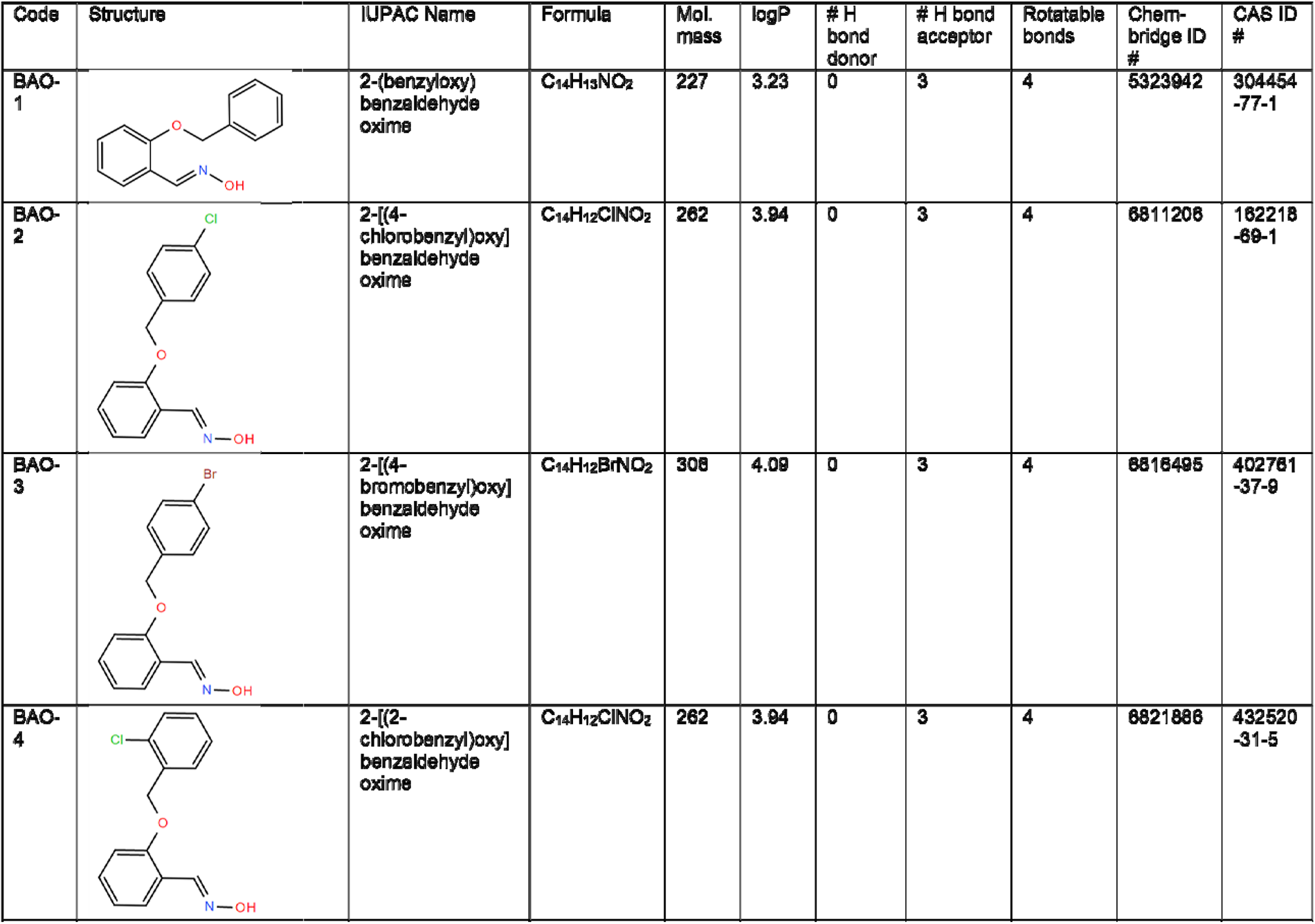
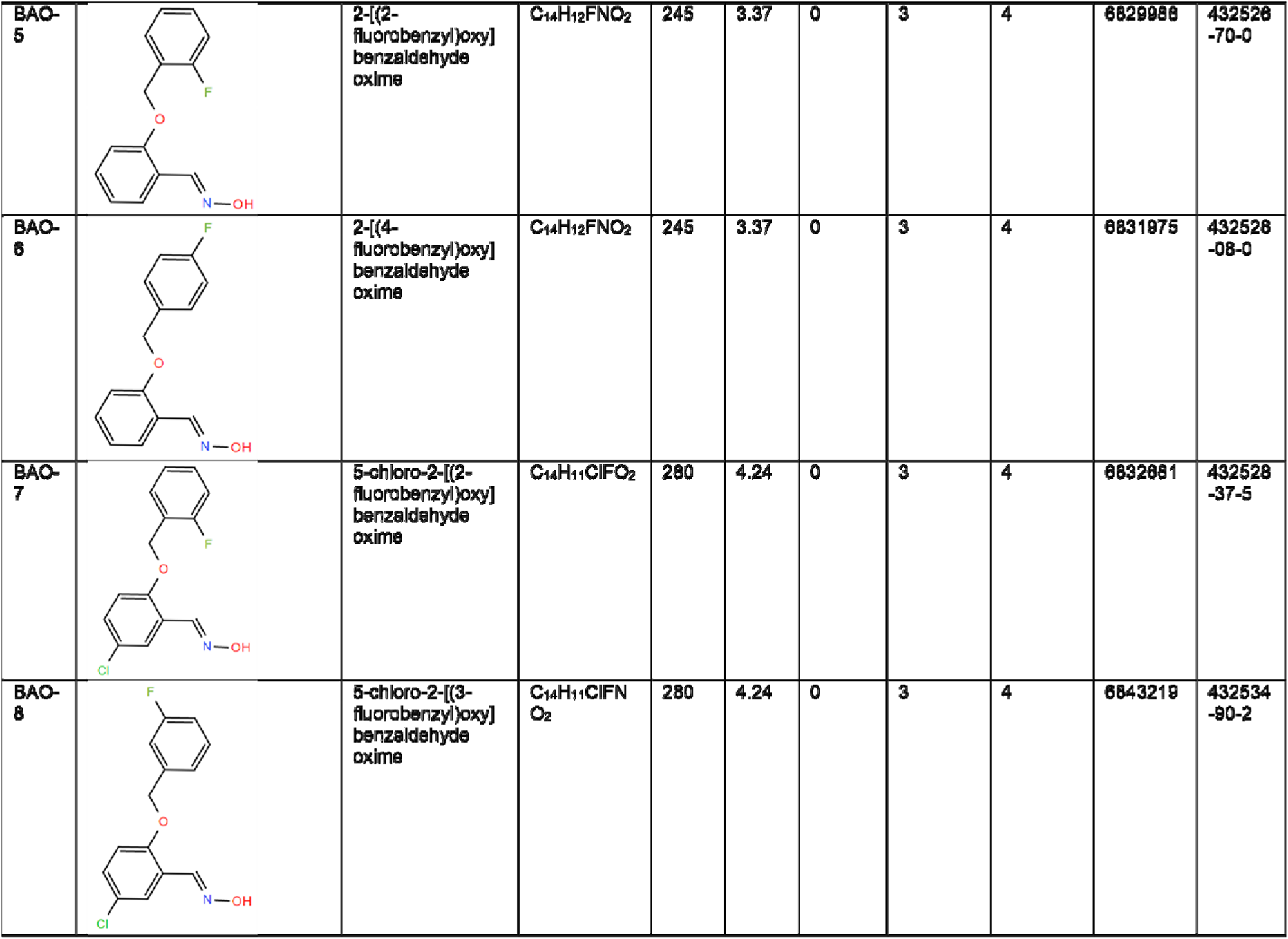
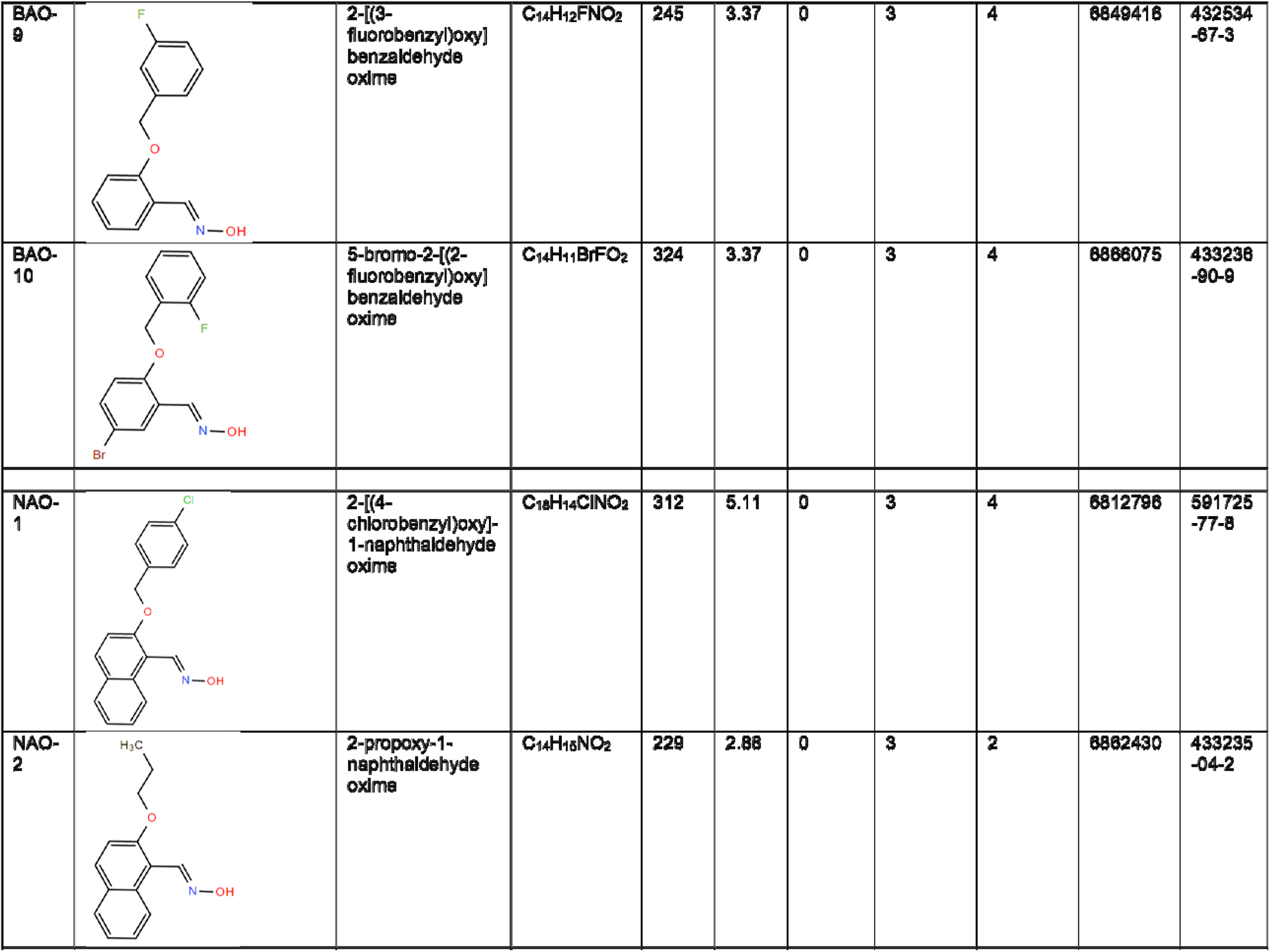
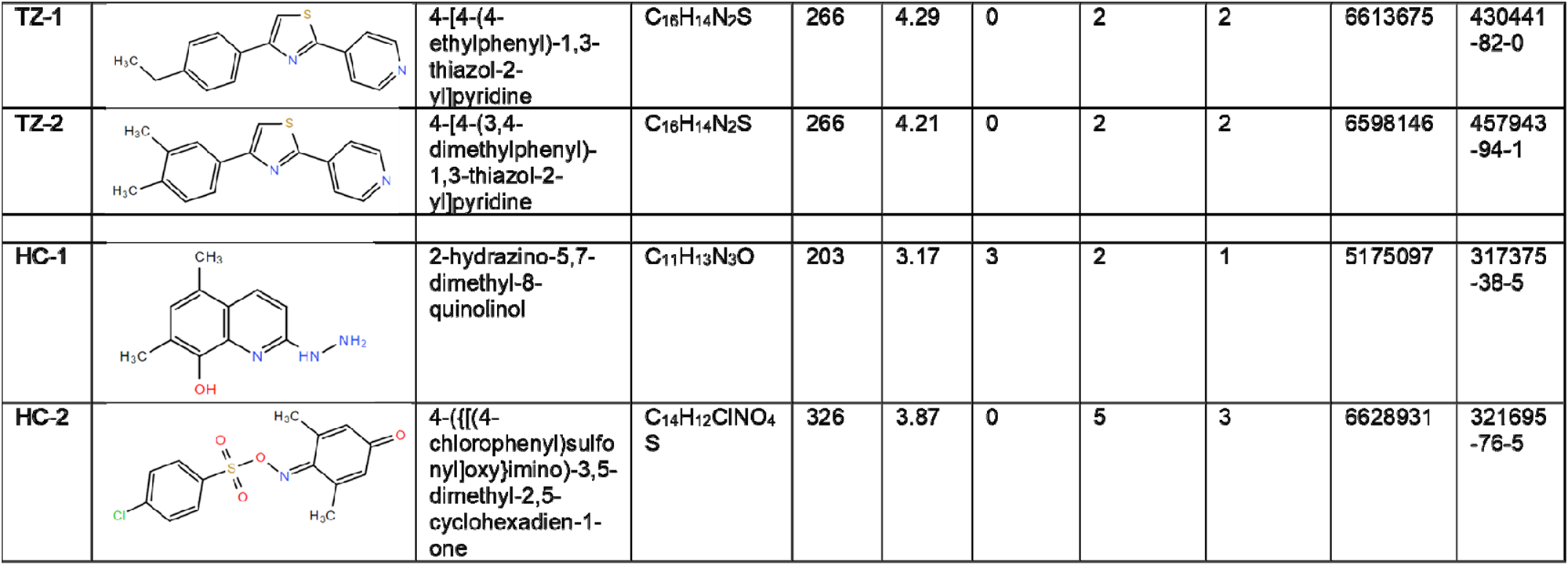
Compounds identified by chemical screening and additional chemical analogues capable of generating altered anthocyanin accumulation phenotypes against a low-temperature challenge with relevant physiochemical properties and identifier data.

### Dose-response analysis: correlating applied compound concentration with degree of observed phenotypic change, and the fine-tuning possible with subtle changes in priming stimuli

An expanded range of concentrations (5, 10, 25, 50, 100 µM – above, including and below the screening condition of 25 µM) of each of the hit compounds and obtained analogues (BAO, NAO, TZ, and HC) were applied to seedlings using the same liquid media protocol to attempt to correlate compound dose with anthocyanin attenuation response. Total anthocyanins were quantified per starting fresh weight of each treated sample, with each sample consisting of 3-4 pooled wells from the 96-well plates containing 8-10 seedlings each (Fig 2).

Several patterns are observable in the results for each compound. Some compounds did not consistently reduce anthocyanin content induced by low temperature compared with the DMSO control: both of the initial hit compounds annotated HC, and BAO-1. The effect of others showed a near-linear inverse relationship, in which higher concentrations of priming treatment predictably resulted in lower levels of anthocyanin: BAO-8, BAO-10 and TZ-1 follow this pattern. BAO-5 was unique in significantly reducing anthocyanin compared with DMSO at all tested doses, but showing minimal difference in levels between doses, perhaps indicating a threshold for activity below 5 µM. Most treatments showed a pattern indicating a certain threshold dose for substantial reduction of anthocyanin content compared with the control: both NAO compounds, BAO-3, BAO-4 and BAO-7 showed a sharp reduction in total anthocyanins starting at 25 µM, with some becoming linear thereafter; BAO-2, BAO-6 and BAO-9 required higher concentrations (50-100 µM) for similar sharp, though often smaller reductions in total anthocyanin accumulation. The concentration range tested did not allow observation of a complete dose-response curve for most compounds (with effect plateaus at high and low doses); approximate least-squares curves were fitted using DMSO control A/g values as dose-zero maxima and zero as hypothetical A/g minima to approximate EC50 values presented in Table 2.

Discernment of molecular features suggesting a specific relationship between structure and efficacy to alter the anthocyanin-attenuation phenotype was not clear at this stage, despite the number of analogues for the BAO category. BAO-1 was ineffective in affecting total anthocyanin content in this assay, with its structure consisting solely of the benzaldehyde oxime skeleton shared by all BAO compounds without additional halogen (F, Cl, Br) additions; this lack of effect may thus suggest significance of the halogen substitutions in triggering the observed phenotypic change. The position of halogen substitution, rather than the specific atom substituted, offered the most insight into functional activity. For example, BAO-5, BAO-6, and BAO-9 showed divergent degrees of effect (approximate EC50 of <5, ∼10, and 50-100 µM respectively) despite each having a single fluorine substitution on the non oxime benzyl ring (in ortho, para, and meta position respectively relative to the oxide bridge). BAO-2 and BAO-4, each with single chlorine substitutions, showed a similar pattern to their fluorine analogues, with the ortho-substituted BAO-4 reducing pigmentation to a higher degree than para-substituted BAO 2 (EC50 ∼5 µM vs. 50-100 µM). Observed here, these variations primarily reveal the exquisite sensitivity of plant systems in perceiving and responding to external cues. Despite the subtlety of the structural differences between the various BAO chemical priming stimuli, the *A. thaliana* seedlings were observed to respond with a fine-tuned alteration of the expected anthocyanin accumulation induced by low temperature exposure.

**Fig 2:**
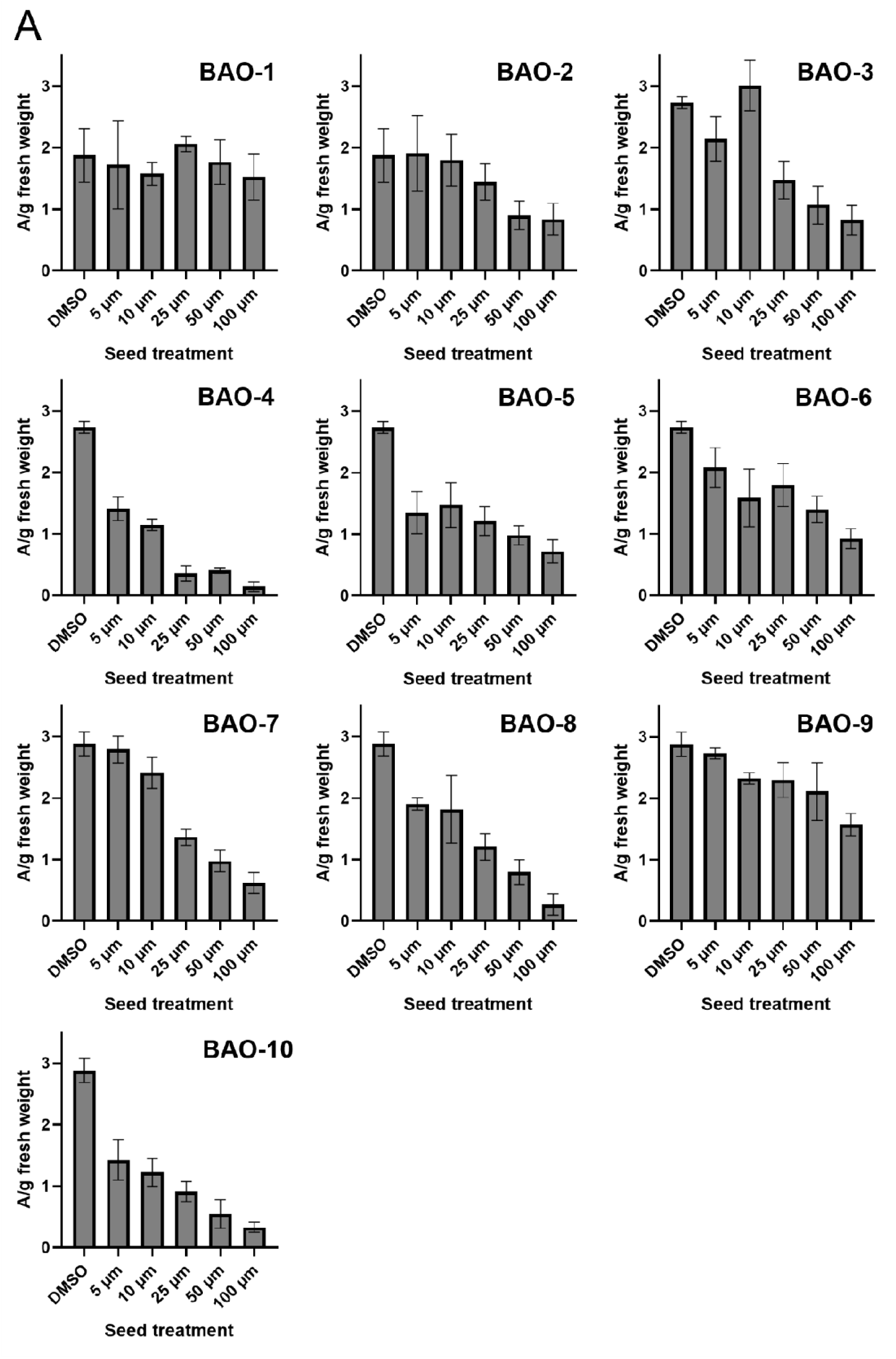
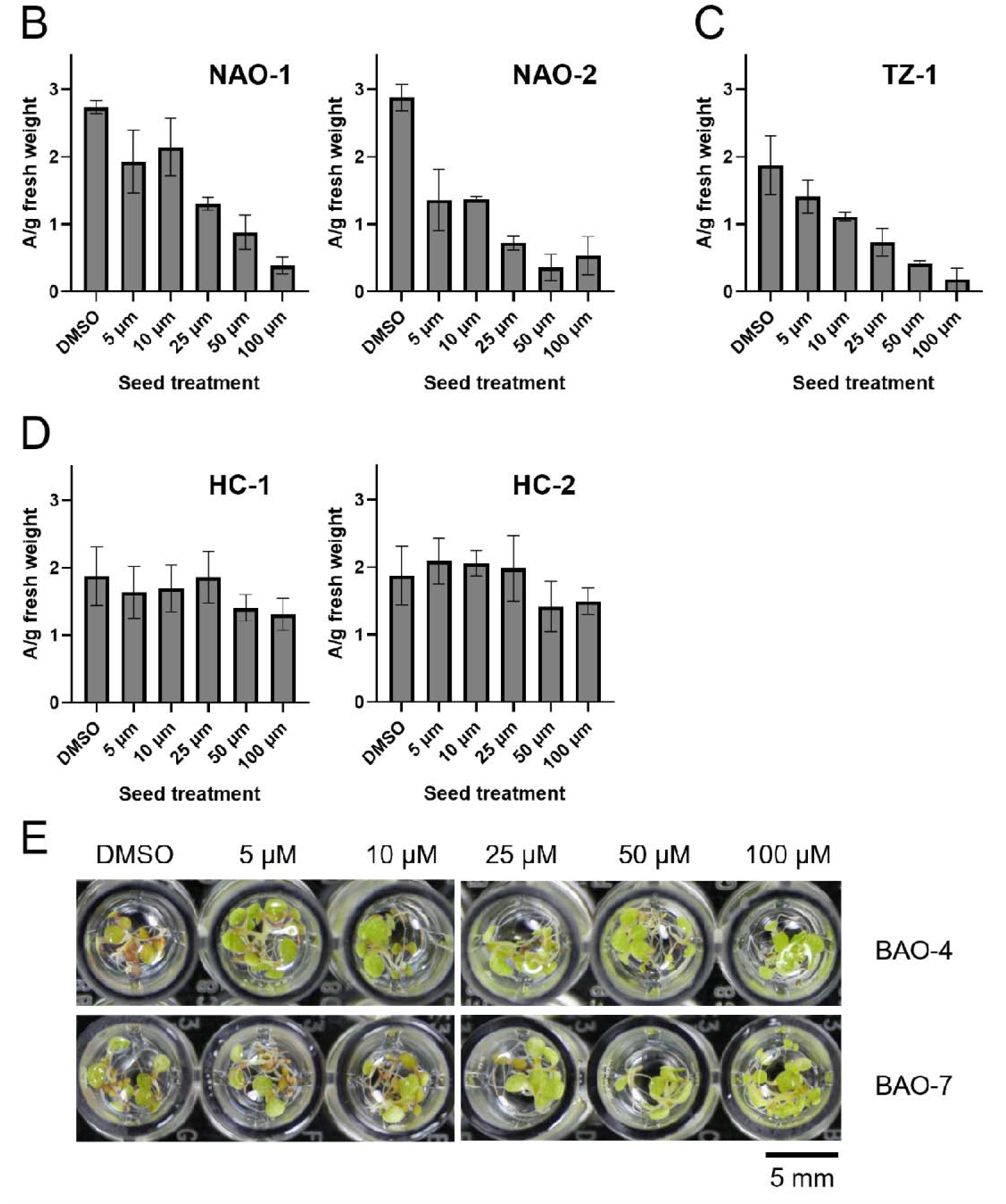
Dose-response analysis shows correlation between anthocyanin attenuation after low temperature and treatment with three structural categories of novel screening compounds. Total anthocyanins assessed by measuring absorbance of extracts divided by fresh mass of each pooled sample (A/g), after treatment with each compound at the concentrations shown (5, 10, 25, 50, 100 µM) for 7 days during stratification and germination, 5 days recovery, then 10 days exposure to 1°C. (A) benzaldehyde oxime (BAO) compounds; (B) naphthaldehyde oxime (NAO) compounds; (C) thiazole (TZ); (D) other hit compounds (HC); (E) example qualitative dose gradient in-plate for two BAO compounds carried forward. Error bars ± SD, n = 2 pooled samples of 20-30 seedlings each.

**Table 2:**
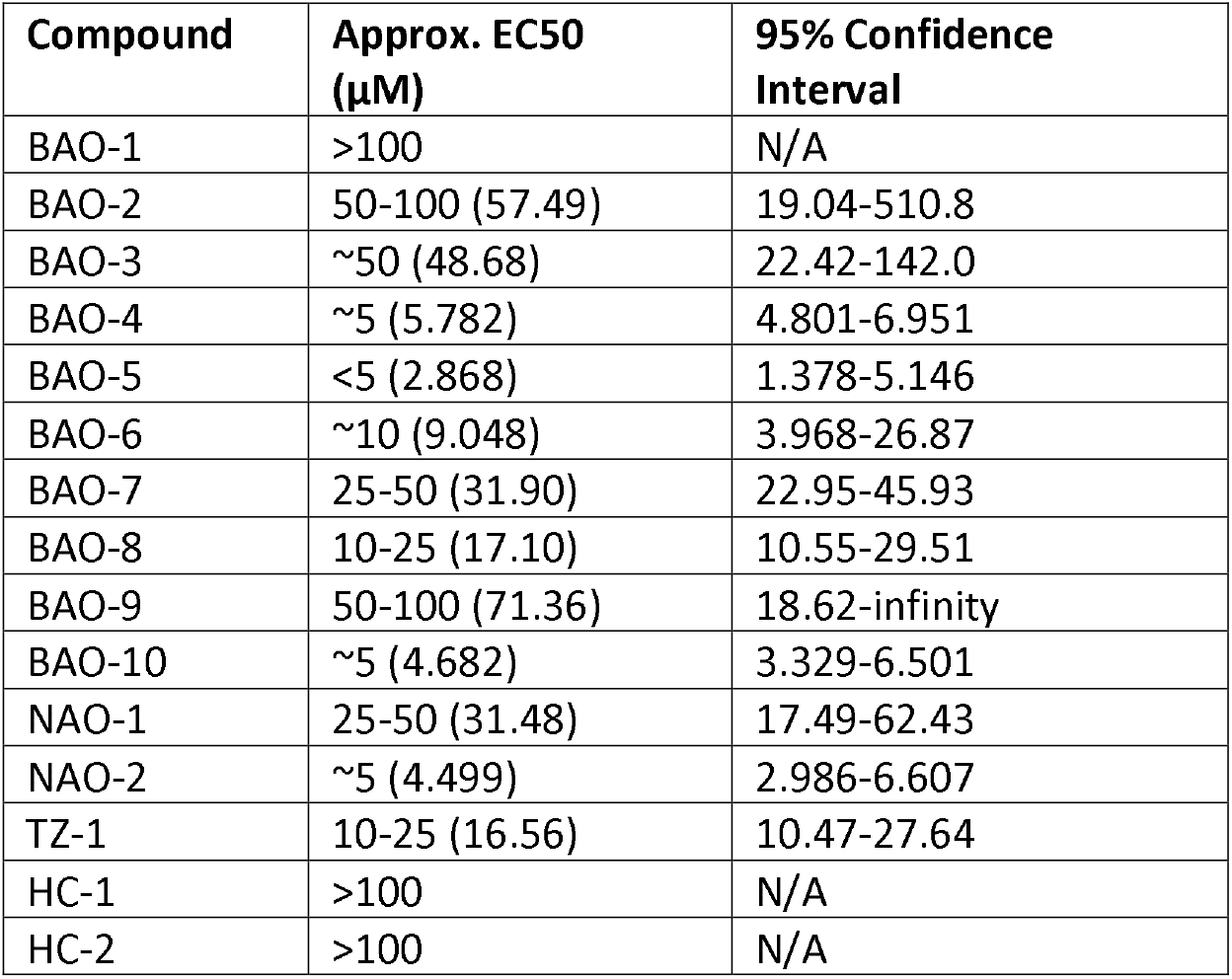
Approximate EC50 values extrapolated from initial dose-response experimental data for BAO, NAO, TZ and other hit compounds (HC) (with calculated value in brackets). For data sets which did not permit curve fitting, no confidence interval was calculated (N/A results).

BAO-4 was chosen from the list of BAO treatments for its high degree of anthocyanin attenuation at the original treatment concentration at 25 µM, and low overall EC50 value (∼5 µM). BAO 7 was carried forward as a contrast to BAO-4, as a dose of 25 µM was sufficient to clearly reduce pigmentation, but by a smaller margin than that induced by BAO-4 and with lesser efficacy to attenuate anthocyanins overall as indicated by EC50. Both NAO compounds were continued. An additional analogue for compound TZ-1 was obtained at this stage due to the strong correlation between compound dose and anthocyanin attenuation for this compound. Three structural categories of chemical priming treatments each with two representative compounds were therefore used in evaluation of the potential energetic or long-term costs of priming with these compounds.

### Chemical priming treatments can generate persistent effects on phenotype through an individual plant’s lifetime at minimal cost to growth and fitness

To evaluate the possible persistence of the identified chemical priming treatments over longer time scales and later low-temperature exposure, an experiment was designed in which seedlings were exposed to chemical priming treatments in multi-well plates as described previously, then transferred to standard potting soil for further growth. After a period of ten days to allow establishment in soil, half the plants were exposed to a brief low-temperature challenge, then grown undisturbed until senescence. The plants in the four experimental combination categories ([chemical priming treatment | DMSO control] x [low temperature challenge | room temperature control]) were observed for alterations in expected vegetative and reproductive development at several representative checkpoints [67]. Anthocyanin accumulation was also assessed after low temperature challenge in one experimental replicate but was not the primary focus of this assay. DMSO control plants did not qualitatively accumulate anthocyanins in response to the low temperature conditions applied at this developmental stage in this growth environment; quantitative analysis revealed a small but not significant degree of induction, with similar results and high standard deviation observed for each chemical priming treatment.

The earliest characteristic in which chemically primed seedlings showed a notable difference from DMSO control treatment was the initiation of inflorescence growth, signalling the beginning of reproductive development (Fig 3A). Seedlings which received certain chemical priming treatments were delayed in reaching this transition, with a lower percentage of individuals achieving this initiation by day 20 of growth vs. DMSO controls. This effect was small for some treatments, with 10-25% of room temperature control seedlings exposed to each NAO treatment delayed, or more extreme, with >80% of seedlings exposed to each TZ treatment in both temperature conditions delayed. This delay was partially recovered by day 23 of growth by all treatment groups and entirely recovered with 100% of plants entering reproductive development by day 24. Exposure to an additional low temperature stimulus delayed this transition for some seedlings in most primed and DMSO sets relative to the control temperature set, with the exception of seedlings exposed to NAO-1: 100% of individuals displayed a bolt bud at day 20, up to four days earlier than other treatment conditions with or without additional stress. Thus, early chemical priming treatments can be seen to effect timing of a developmental process beyond germination and early seedling development, possibly integrating response to later environmental conditions, with the degree and pattern of effect unique to each treatment.

Several growth parameters spanning other aspects of vegetative and reproductive development, specifically final rosette diameter, dry above-soil biomass at senescence, and inflorescence length were assessed (Fig 3B,C; Fig S3A). Chemical priming treatment alone or combined with subsequent low temperature treatment either did not affect final rosette diameter (both BAO treatments) or resulted in reductions in growth for some combinations (NAO-1 with chilling; NAO-2 without chilling; both TZ treatments without chilling). Although these reductions in diameter were statistically significant (p < 0.05), the magnitude of the change was small (Fig 3B). Plants treated with BAO and TZ compounds showed negligible difference in inflorescence growth with or without additional chilling exposure, in contrast to the lag in initiation for TZ treated seedlings especially. The treatment group showing the greatest lag in inflorescence growth, NAO-2 primed plants exposed to subsequent low-temperature, nevertheless was statistically equivalent to DMSO controls by the end of the observation period (Fig S3A). Differences in total dry biomass after senescence were observed for all three chemical priming treatment groups (Fig 3C). Plants primed with BAO-4 were lower in total mass than DMSO controls both with and without subsequent low temperature challenge while plants primed with BAO-7 were lowest in mass in the control temperature, but not significantly different from controls if exposed to cold. Plants primed with NAO-1 exhibited significantly higher biomass than DMSO in control temperature, but significantly lower with the additional low temperature challenge; plants primed with NAO-2 were also significantly lower in mass when exposed to the second stress. Plants primed with both TZ compounds showed significantly increased biomass in control temperature, perhaps surprisingly due to their delayed start in inflorescence growth and smaller rosette diameter; plants primed with TZ-1 with additional low temperature challenge were also significantly greater in biomass compared with control plants that received DMSO.

**Fig 3:**
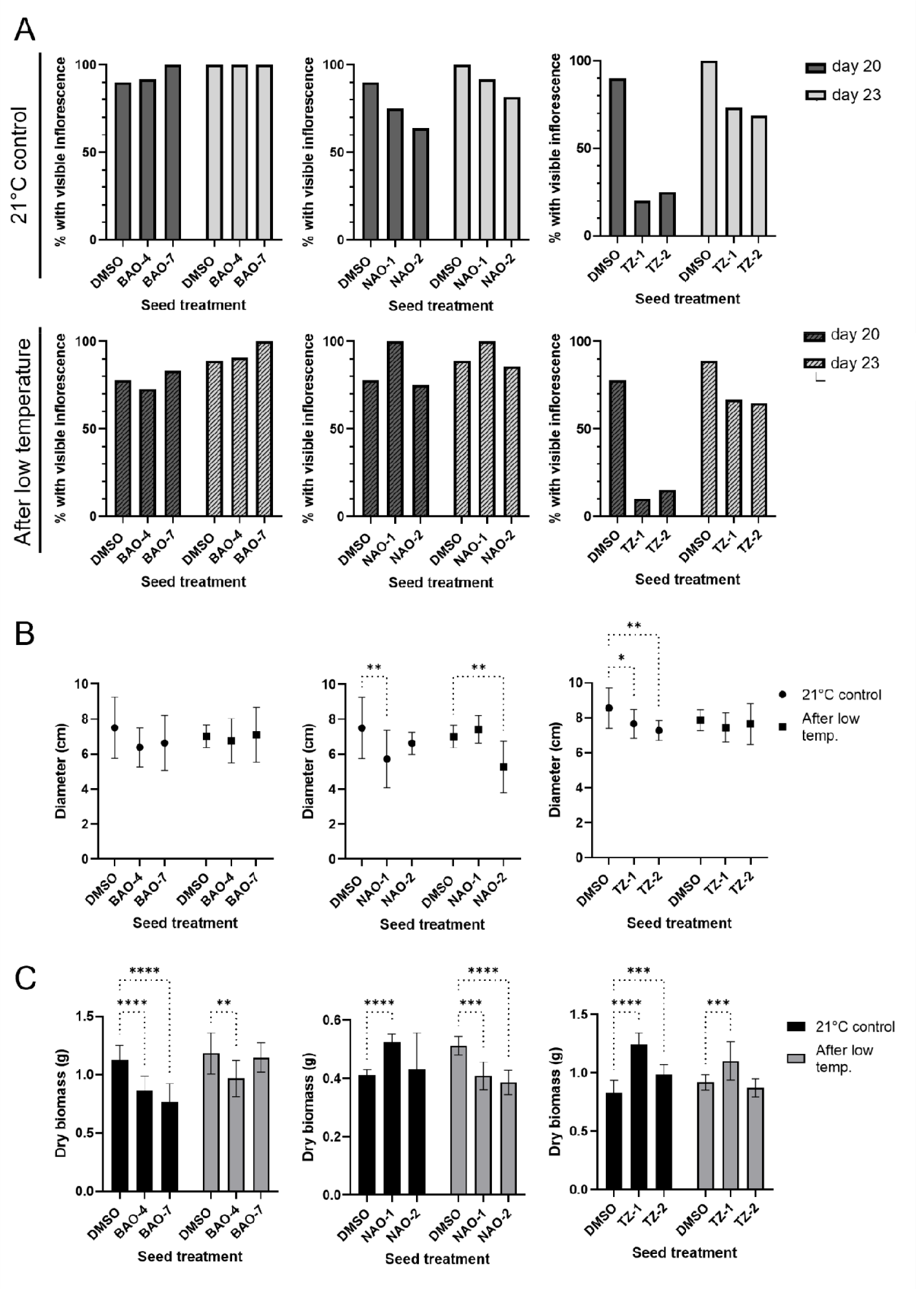
Morphometric analysis of developmental and reproductive characteristics of seedlings grown to senescence after chemical seed priming does not indicate inherent ‘cost’ due to early priming treatment. (A) Percentage of plants entering floral transition at days 20 and 23 of growth of primed and control plants, without (top) and with subsequent low temperature challenge (bottom), assessed by emergence of visible inflorescence. n ≥ 8 for all treatment combinations. (B) Rosette diameter observed during week 5 of growth (day 37 for BAO, NAO; day 39 for TZ). n ≥ 8 for all treatment combinations, error bars ± SD. Significant difference tested with two-way ANOVA (p < 0.05 (*), p < 0.01(**)). (C) Dry above-ground biomass after senescence. n ≥ 8 for all treatment combinations, error bars ± SD. Significant difference tested with two-way ANOVA (p < 0.05 (*), p < 0.01(**), p < 0.001(***), p < 0.001(****)).

Seeds harvested from plants primed with each of the chemical priming treatments and from each chilling exposure group were harvested and assessed for viability (Fig S3B). Near 100% germination rates (with some variability) were observed for all treatments, with the exception of one control DMSO treatment (no chilling, grown alongside plants primed with NAOs). The strong germination rates displayed otherwise and by the DMSO results in the other panels at this time point suggest that early seed priming does not typically affect seed viability in the resulting plants. Whether any effects persist from seed priming in the parent generation into later growth stages of S1 plants was not consistently established.

The alterations in growth patterning induced by chemical priming considered alongside their previously observed effect to reduce anthocyanin accumulation induced by low temperature generated some apparent contradictions. A treatment which causes a plant to accumulate lower levels of anthocyanins may indicate that the plant was less perturbed by an inducting stress, but that plant may also be more susceptible to later challenge as anthocyanins have protective characteristics [49,52,72]. The biomass data of NAO-1 treated plants support this latter hypothetical relationship, as unchallenged plants show enhanced growth, perhaps due to reduced metabolic energy expenditure on flavonoids, but reduced growth when challenged with cold. However, BAO-4 treated plants show similar biomass accumulation with or without cold challenge, as do those treated with TZ-1. Additionally, while both BAO-4 and TZ-1 significantly attenuate anthocyanins in the wellplate system, their effect on total biomass diverges, with BAO-4 plant mass consistently reduced compared with DMSO, and TZ-1 increased. This divergence suggests the possibility of different overall mechanisms of effect between the structural categories, de-emphasizes the importance of the response to low-temperature challenge in the effect of the compounds, and resists an overall generalization of anthocyanin regulation and stress responsiveness in the chemically primed plants.

These data together show that the effects of chemical priming with the novel BAO, NAO and TZ compounds identified can persist throughout the lifespan of the treated plant. The specific effects of each treatment regarding a trait of interest should be assessed and optimized individually, as significant variation can be seen in effect between highly similar chemical analogues. The concept of “costs” or growth-defense trade-off for each treatment should also be assessed individually, and according to specific traits: TZ treatments, for example, delayed the onset of floral transition, but recovered and generated increased total biomass at the time of senescence. Altogether, the small degree of phenotypic alteration, recovery of delays, and lack of effect on seed viability suggest that there is minimal inhererent cost to chemical priming with novel agents as an overall approach.

### Chemical priming treatment can be reduced in duration, and effect differs based on specific phase of application

Germinating seeds are understood to undergo a three-phase process. Phase I consists primarily of initial water uptake by the seed. This triggers phase II, which is recognized by a lag in water uptake but the activation of cellular and metabolic processes including DNA repair, cell cycle activation and protein synthesis. The changes of phase II produce the conditions necessary for phase III, cell elongation and weakening of the seed coat allowing emergence of the radicle, resumed water uptake, and further growth [29,73]. The screening protocol used to identify the BAO, NAO, and TZ compounds as chemical priming agents applied the compounds to seeds for a period of 7 days – 4 days stratification at 4°C in complete darkness, permitting imbibition but inhibiting further germination, then a further 3 days at 21°C on a long-day photoperiod. This treatment period fully encompasses the time until radicle emergence and full germination, typically observed by the third day at 21°C in light. With thought to potential future usefulness in the field, and to reflect the different processes occurring at different phases of germination, we sought to reduce the number of days of chemical priming treatment and identify a possible key application time window related to efficacy in altering later phenotype. The chemical priming regime was therefore applied for a reduced window of 3 days, either during dark stratification or light germination, and anthocyanin attenuation assessed before and after subsequent chilling exposure for plants treated with BAO and NAO analogues (Fig 4A-B). The duration of 3 days was applied to both phases for consistency and was chosen due to being the shorter of the two original phases (dark-stratification vs light-germination). This choice also preserved the minimum recovery period of 5 days for the light-germination treatment group prior to the low temperature challenge (with original stress onset maintained) to keep the duration of possible effect persistence consistent.

Seedlings treated with NAO compounds showed significant reductions in anthocyanin content with both three-day chemical priming regimes; NAO-1 only after further chilling treatment, and NAO-2 both before and after additional chilling exposure. Although significantly reduced in both assays, the degree of attenuation by NAO treatment was greater in those plants treated during germination. Seedlings treated with the two BAO compounds showed divergent results, where both priming compounds and three-day priming regimes significantly reduced anthocyanin content prior to chilling. However, after chilling BAO-4 significantly reduced anthocyanins in both regimes, but plants treated with BAO-7 during light-exposed germination produced reduced levels of pigment, where plants treated with BAO-7 only during dark stratification produced comparable anthocyanin levels to DMSO control plants. The heterogeneity of these results, especially the complete loss of effectiveness of BAO-7, suggests that despite their similar capacity to cause anthocyanin attenuation under the initial screening protocol, different regulatory mechanisms may be at play in their biological effect. Possible explanations for these differences may be the variance in their activity as treatments (Table 2) but may also lie in a requirement for priming compound presence during a certain germination process (e.g. the DNA repair processes in Phase II), or perception and/or signaling related to light exposure.

An additional experimental protocol further reducing chemical priming duration to a single day was run concurrently with the three-day application regimes for each individual day within the three day lighted germination period (Fig 5). For all combinations of priming treatment and chilling exposure, seedlings treated with DMSO or chemical priming compounds on either day 1, day 2, or day 3 of growth conditions produced levels of anthocyanin accumulation which were not statistically significantly different from the other single-day exposures of the same treatment. The two BAO priming treatments showed inverse significance when applied for one day: BAO-4 did not significantly reduce total anthocyanin compared with controls prior to chilling for two of the three tested single days, but did significantly reduce levels induced after later chilling; BAO-7 significantly reduced pre-chilling anthocyanin levels, but this attenuation was not significant when observed after chilling. NAO-2-primed seedlings showed significant attenuation compared with DMSO across all but one treatment period in both time points; seedlings primed with NAO-1 did not achieve significantly lower levels of anthocyanin prior to chilling treatment, but after further chilling stress exposure and anthocyanin induction, NAO-1-primed seedlings retained pre-chilling total anthocyanin which was significantly lower than that induced by chilling in DMSO-treated controls. Together, these results demonstrate that one day of chemical seed priming during light-exposed germination can be sufficient to induce persistent phenotypic effect, but that this is not universally true for all compound treatments. Given the differences observed within this small number of priming treatments, it seems likely that the key treatment window and minimum treatment duration would require case-by-case optimization for each proposed chemical priming agent, but the possibility of lasting phenotypic effects resulting from a single day of low-dose priming remains promising.

**Fig 4:**
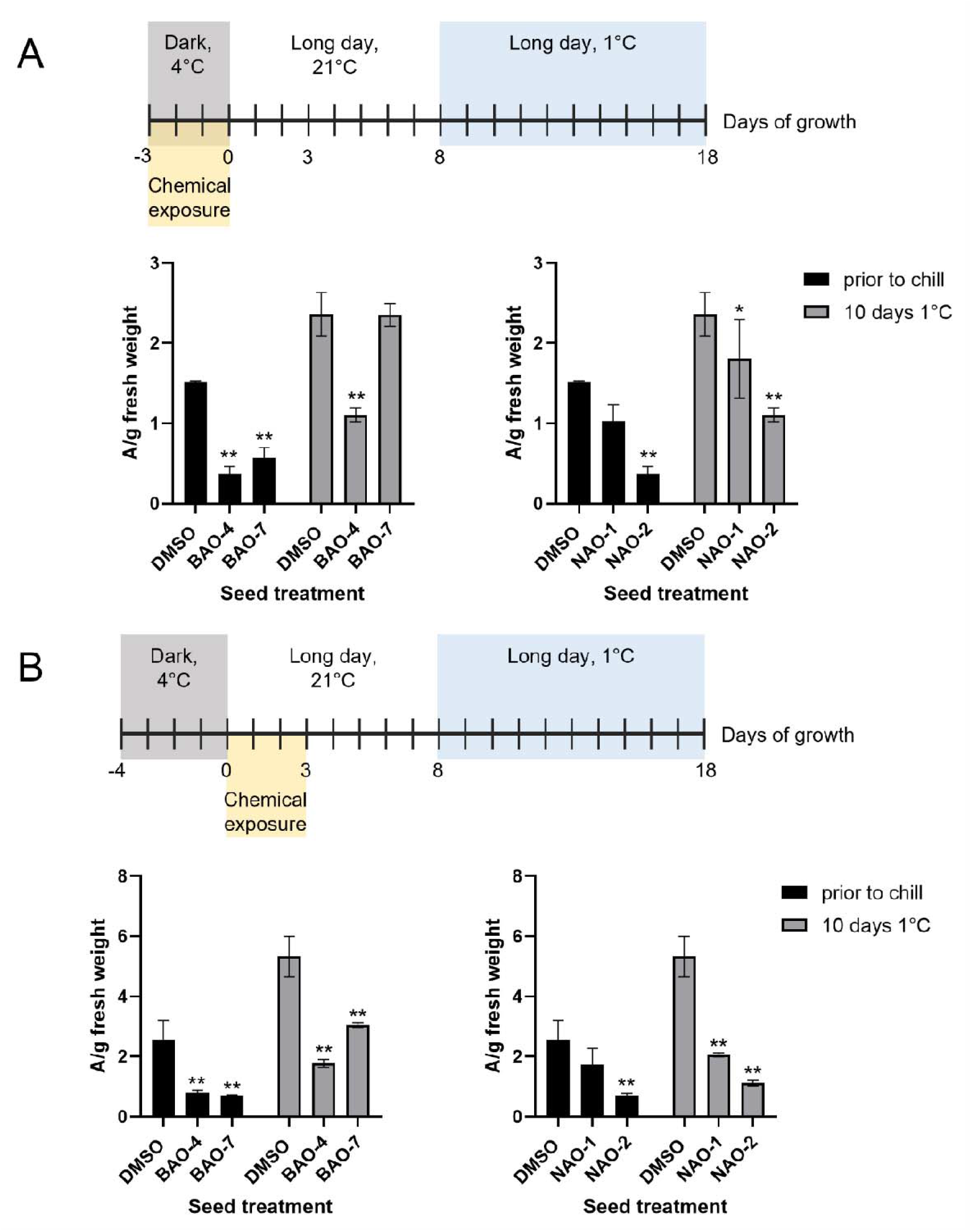
Reduced priming duration of 3 days during either dark stratification or light germination revealed light-grown germination as preferred window for treatment. BAO and NAO priming compounds were applied either during dark-grown stratification (A) or light-grown germination (B); anthocyanin content before and after low temperature shown. Error bars ± SD, n = 3 pooled samples of 20-30 seedlings each. Significance assessed with two-way ANOVA against DMSO control in each group, Dunnett’s multiple testing correction, p<0.05 (*) and p<0.001 (**) indicated.

**Fig 5:**
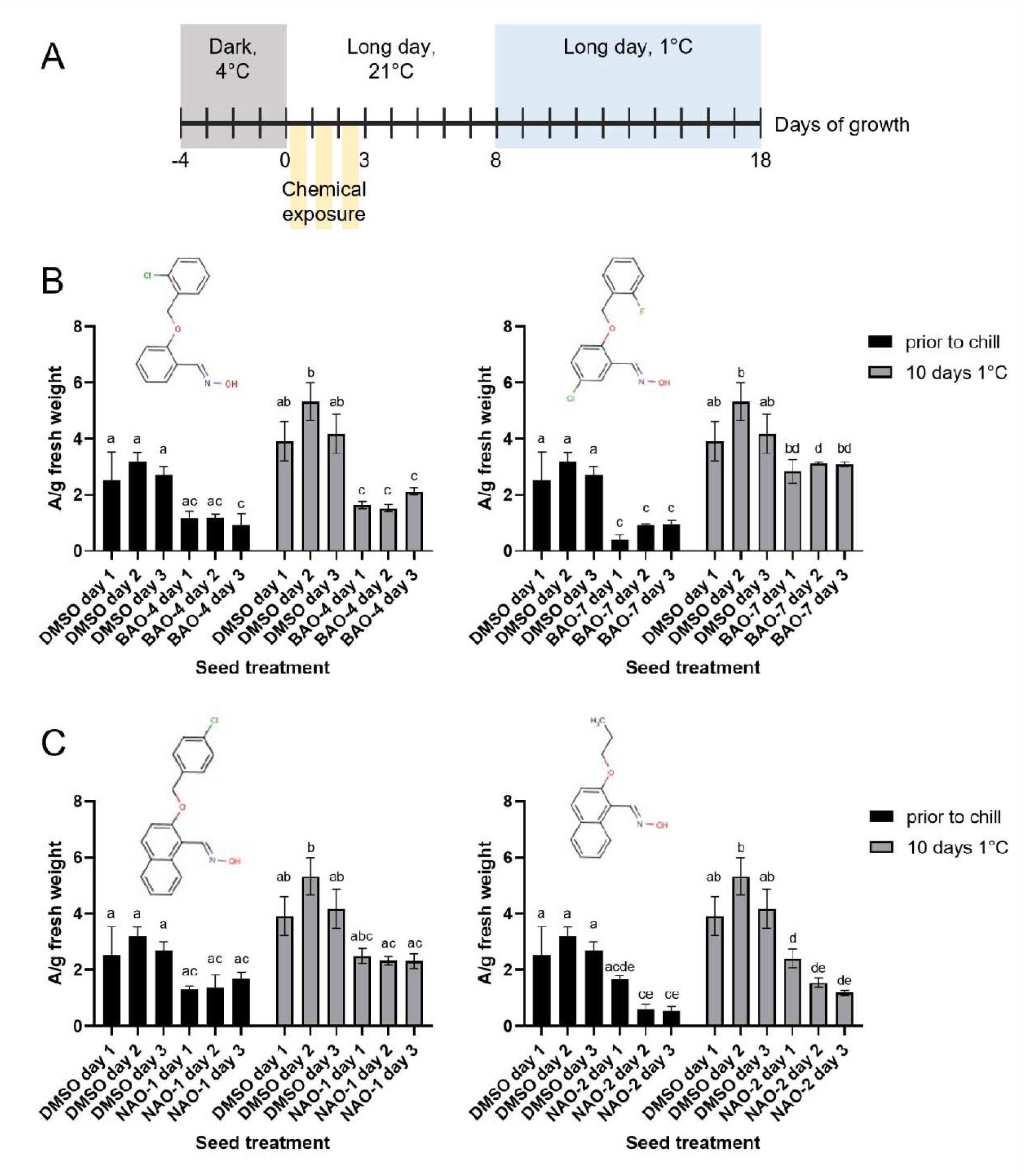
One day of compound exposure during light-grown germination can be sufficient to prime for anthocyanin attenuation before or after subsequent low temperature challenge, dependent on treatment. Anthocyanin attenuation vs. DMSO controls shown for treatment on each of three possible single day treatment windows during germination (A); BAO compounds (B); NAO compounds (C). Error bars ± SD, n = 3 pooled samples of 20-30 seedlings each. Significance assessed with two-way ANOVA comparing all treatments with all other treatments, Tukey’s multiple testing correction, p<0.05. Results which were not significantly different are annotated with the same letter.

Although the original screening assay observed qualitatively visible anthocyanin accumulation induced by low temperature, the additional quantitative data in these modified experiments revealed that sufficient anthocyanin levels were also present in the plants prior to stress challenge to permit statistically significant attenuation by chemical priming. Wild-type *A. thaliana* plants accumulate flavonoids and anthocyanins in an age- and tissue-dependent manner, with accumulation noted at the junction of the hypocotyl and cotyledons of young seedlings which decreases over time [56,74]. Given the developmental stage of seedlings at day 8 of growth in the chemical priming assay, this is consistent with detectable anthocyanin levels prior to additional induction, but does suggest that the attenuation effect of chemical priming treatment is not strictly connected to the secondary cold challenge. This possibility that the effect of chemical priming primarily related to anthocyanin regulation specifically was further explored using the BAO category of compounds.

### Beyond a single abiotic stimulus: effect of small molecule seed priming to attenuate anthocyanin accumulation induced by different treatments

In parallel with the chemical library using low temperature stimulus in the presence of sucrose, a similar screen to identify small molecule priming treatments capable of altering plant response to nitrogen deprivation (Naik, 2016) was developed (Fig 6). This screen utilized liquid MS growth media supplemented with ammonium nitrate (NH_4_NO_3_) and sucrose to achieve a “moderately high” C:N ratio of 5:1, which was demonstrated to stimulate primary root growth, chlorophyll breakdown and anthocyanin accumulation consistent with an expected nitrogen-deprivation phenotype in wild-type seedlings, without excessive bleaching or growth inhibition. This experimental protocol was used to assess cross-stress effects of BAO chemical priming to better understand if the activity of these compounds related to anthocyanin attenuation was specific to low-temperature response, or a more general effect on this metabolic pathway.

In the nitrogen-deprivation assay, seedlings in the nitrogen-sufficient control condition (N+) accumulated relatively low levels of anthocyanins in all treatment categories with the lowest levels observable in plants treated with both BAOs . When nitrogen was depleted in the growth media, both BAO analogues significantly attenuated anthocyanin accumulation compared with controls (Fig 5B). These results confirm the ability of the BAO compounds to act as chemical priming treatments to alter phenotypic response to abiotic stimuli after their removal. The results of this assay further suggest that the activity of this structural category of molecules is not specific to the environmental condition, low temperature, that was used to identify it.

Anthocyanin regulation by both temperature and nutrient deprivation signals has been characterized in detail in the literature. Low-temperature induction of anthocyanin biosynthesis is known to be light-dependent and regulated by the HY5-COP1 module, in which the repressive COP1 degrades HY5 when both are localized to the nucleus in the dark; in light, COP1 is re-localized to the cytosol allowing HY5 to upregulate anthocyanin production through multiple mechanisms and interaction partners. Cold temperature also triggers COP1 re-localization and similarly releases repression of anthocyanin biosynthesis [60,75,76]. In response to nitrogen availability, an entirely distinct gene family regulates anthocyanin accumulation: the *LATERAL ORGAN BOUNDARY DOMAIN* (LBD) transcription factor gene family paralogs *LBD37, LBD38* and *LBD39* repress transcription of anthocyanin biosynthetic regulators *PAP1* and *PAP2* when nitrogen is available, and release this repression allowing for pigment accumulation upon onset of nitrogen deficiency [60,77]. Given the flexibility of the chemical priming treatments to attenuate anthocyanin induction by both low temperature and low nitrogen and the distinct regulatory modules related to each of these environmental signals, these results do not suggest that either the HY5-COP1 module nor the LBD module are likely to be direct targets of chemical priming activity. The lack of specificity regarding abiotic stimulus may therefore instead reflect that the chemical priming treatment activity is related to a more general stress-related plant response, such as ROS homeostasis and signaling [78], or direct effect on the regulation of anthocyanin production and maintenance itself.

**Fig 6:**
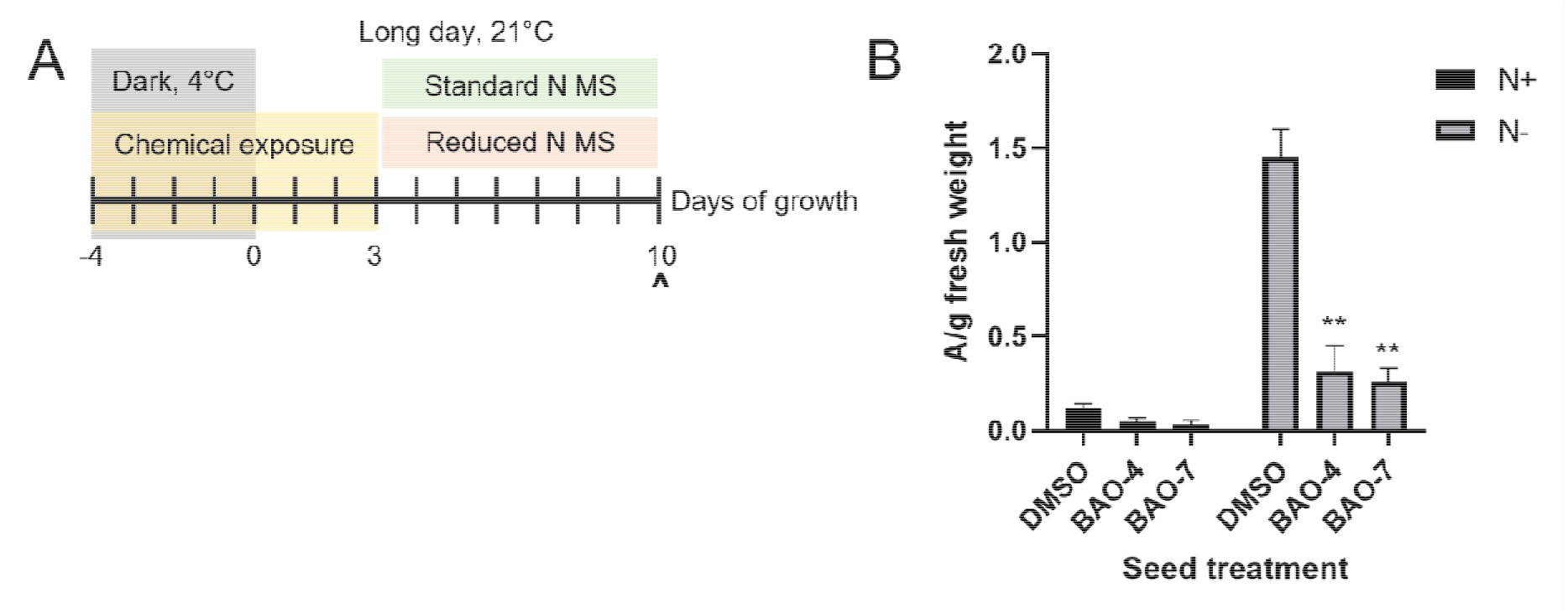
Benzaldehyde oxime chemical priming treatments identified by environmental screening for altered anthocyanin response to low temperature can also attenuate anthocyanin induction by nitrogen deprivation. (A) Experimental timeline for chemical priming and application of nitrogen insufficient media to Col-0 wild-type seeds. (B) Anthocyanin attenuation response of seedlings treated with BAO compounds in the low nitrogen protocol. Error bars ± SD, n = 3 pooled samples of 20-30 seedlings each. Significance assessed with two-way ANOVA, p<0.05 (*) and p<0.001 (**).

To further differentiate the activity of these chemical priming treatments from the secondary environmental challenge conditions used to induce phenotypic responses of primed plants, a mutant line which constitutively overproduces anthocyanins and other flavonoids was used (Fig 7). The *production of anthocyanin pigment 1-Dominant (pap1D)* mutant line was originally generated in a screen using a T-DNA activation tagging strategy to identify novel functional genes, in which wild-type *A. thaliana* Col-0 plants were transfected with a vector containing multiple copies of the strong constitutive CaMV-35S promoter element [69]. *pap1D A. thaliana* plants are visibly purple with unusually high levels of anthocyanin pigments produced in all tissues (Fig 7C), as the activation tag in this line inserted adjacent to the gene subsequently identified as PAP1, a R2R3-MYB transcription factor now known to be part of the Myb-bHLH-WD40 complex which upregulates the transcription of multiple flavonoid biosynthetic enzyme genes in *A. thaliana* [46,69]. Transgenic plants overexpressing *PAP1* have previously been used effectively to dissect anthocyanin regulation in response to external cues including temperature, nitrogen, and sucrose [57,79]. *pap1D* plants have also been used as a model to connect transcriptional and metabolic regulation and identify new anthocyanin biosynthetic enzyme genes [80].

In the absence of any further environmental cue, BAO priming treatments persisted after removal to attenuate anthocyanin accumulation in *pap1D* seedlings (Fig 7). Qualitative observation of *pap1D* seedlings at room temperature and wild-type seedlings after low-temperature treatment showed similar intensity of visible anthocyanin colouration with comparable localization in the emerging true leaves and cotyledon margins (Fig 7A-C). A modified chemical priming assay was designed in which BAO and BI seed treatments were applied for a seven-day period through stratification and germination, consistent with earlier experiments (Fig 7D); priming treatments were removed by rinsing, then seedlings were grown with no further perturbation at 21°C for a seven-day recovery period. Chemical priming of *pap1D* seedlings with both BAO treatments resulted in significantly reduced anthocyanin levels in *pap1D* seedlings compared with DMSO control treatments (Fig 7E). Attenuation of anthocyanin accumulation by BAO priming in this context further supports the conclusion that the effect of the chemical priming treatment primarily relates to general anthocyanin regulation, rather than signaling related to perception or response to a specific environmental stress stimulus.

Given that sucrose supplementation of the liquid MS media growth system was necessary to observe any anthocyanin induction, and the well-documented ability of sucrose to induce anthocyanin accumulation in the literature, exogenous sugar treatments were explored as a third stimulus variable to assess the attenuative ability of the chemical priming treatments. High levels of sucrose applied after the removal of the priming compounds were used to assess chemical priming capacity to attenuate anthocyanin induction by a third mechanism. Wild-type seedlings were exposed to the two BAO chemical priming treatments for a seven-day period throughout stratification and germination as described previously, with priming treatments removed by rinsing and seedlings allowed to grow undisturbed at 21°C for five days of recovery, parallel with the low-temperature assay (Fig 8). At 30 mM sucrose, the concentration added to the MS medium used for all prior temperature assays, both DMSO control-treated seedlings and the chemically primed seedlings show negligible anthocyanin content from day 11 onwards, as expected developmentally (Fig 8B; [56,74]). When 100 mM sucrose media was added, seedlings of all treatments accumulated additional anthocyanin by day 14 after a brief delay. While levels were attenuated in seedlings primed with both BAO treatments at days 8 and 11, by day 14 anthocyanin content in the primed seedlings was not significantly different from DMSO and continued to increase similarly at day 17 (Fig 8C). When 200 mM sucrose media was added, anthocyanin content of all seedlings increased dramatically. Both BAO-treated groups increased immediately, while DMSO treated seedlings exhibited a delay until day 14 similarly to the 100 mM treatment. However, by day 17, DMSO-treated plants continued to accumulate additional anthocyanin where levels in the BAO-treated plants had either slowed or plateaued (Fig 8D). The subtlety of effects observed in this assay, in the degree of delay and possible limitation on maximum accumulation differentially revealed by varying sucrose application, suggests a finely-tuned effect of the priming compounds. These patterns show that anthocyanin upregulation mechanisms are not abolished but rather dampened, and can be partially overcome with a sufficiently extreme inductive stimulus like the highest sucrose treatment used here.

**Fig 7.**
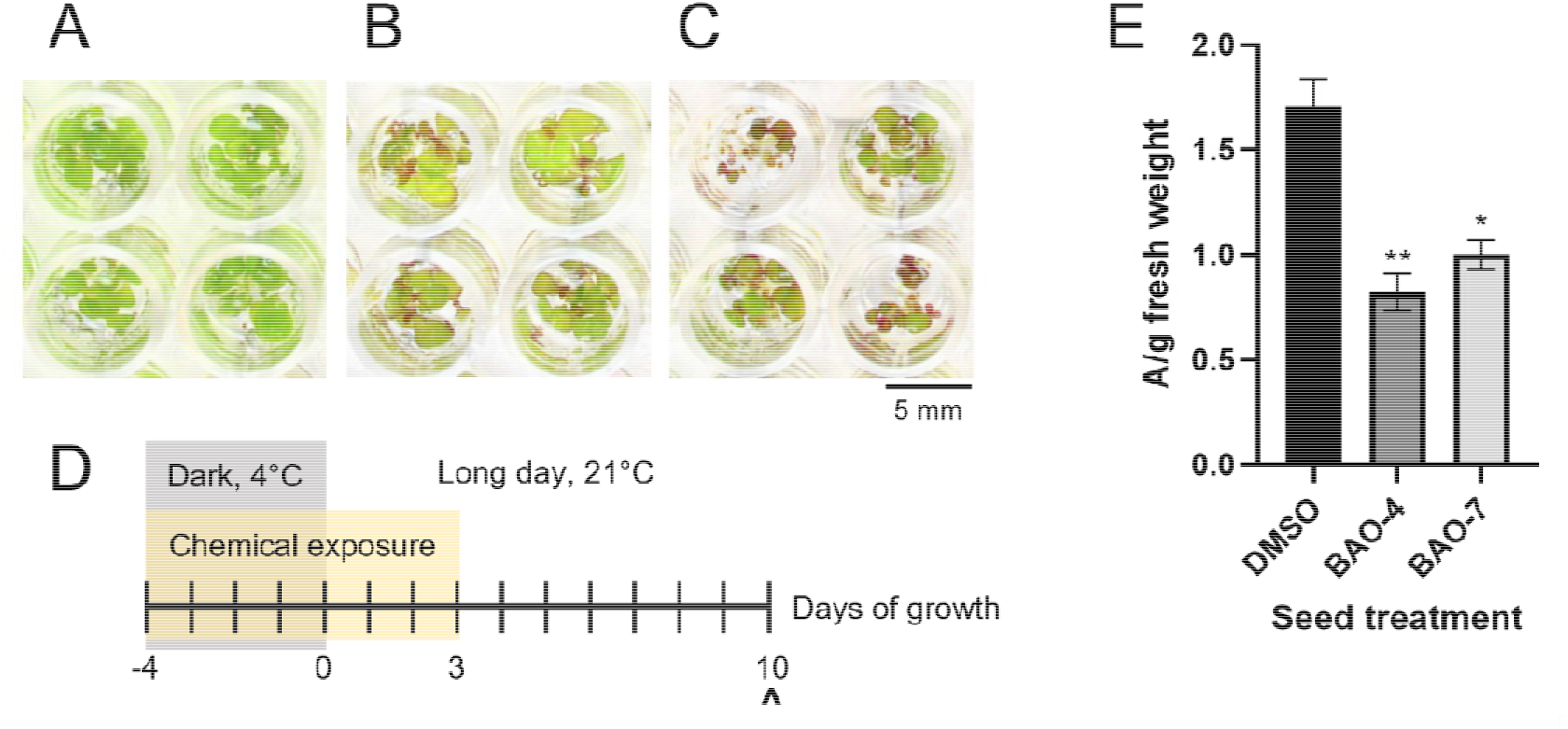
Benzaldehyde oxime chemical priming treatments can suppress endogenous overproduction of anthocyanins in the pap1D overexpression mutant line. (A) Wild-type seedlings at 8 days of growth at 21°C. (B) Wild-type seedlings grown as in A, then exposed to 1°C chill for 10 days. (C) *pap1D* seedlings at 8 days of growth at 21°C. (D) Experimental timeline for chemical priming and subsequent observation (∧) of *pap1D* seedlings after a recovery period. (E) Quantification of anthocyanin content in chemically primed pap1D seedlings 7 days after compound removal. Error bars ± SD, n = 3 pooled samples of 20-30 seedlings each. Significant difference vs. DMSO assessed with one-way ANOVA shown, p<0.05 (*) and p<0.001 (**).

### Overall conclusions

Although both seed priming and chemical genetics are well-developed strategies within the plant research literature, this work represents a novel combination of the two fields of inquiry. This effort to uncover bioactive small molecules which may be applied to seeds at low concentration, removed, and still alter a biological relevant characteristic such as anthocyanin accumulation successfully identified several structural categories of molecules for further inquiry. The BAO, NAO and TZ compounds here attenuate anthocyanins in a dose-dependent manner and do not cause major growth impediments in plants grown to senescence after early priming. Although the precise target of the priming treatments was not identified in this work, the flexibility of the treatments to attenuate anthocyanins induced by different exogenous and endogenous mechanisms suggests that their mode of action is likely to regulate some aspect of the synthesis, stability or possibly degradation of these metabolites. The structural diversity between the identified priming molecule categories, the different degree of activity of each structural category and nuances such as differences in senescent biomass may suggest different target pathways for each. The biochemical mechanisms by which information from prior life experience are retained in plants in priming phenomena are still yet to be comprehensively understood. Small molecules persistently affecting a metabolite class relevant to both plant development and stress response may be a useful tool, more specific than the broad-spectrum changes induced by natural abiotic treatments, for teasing apart the regulatory mechanisms involved.

**Fig 8.**
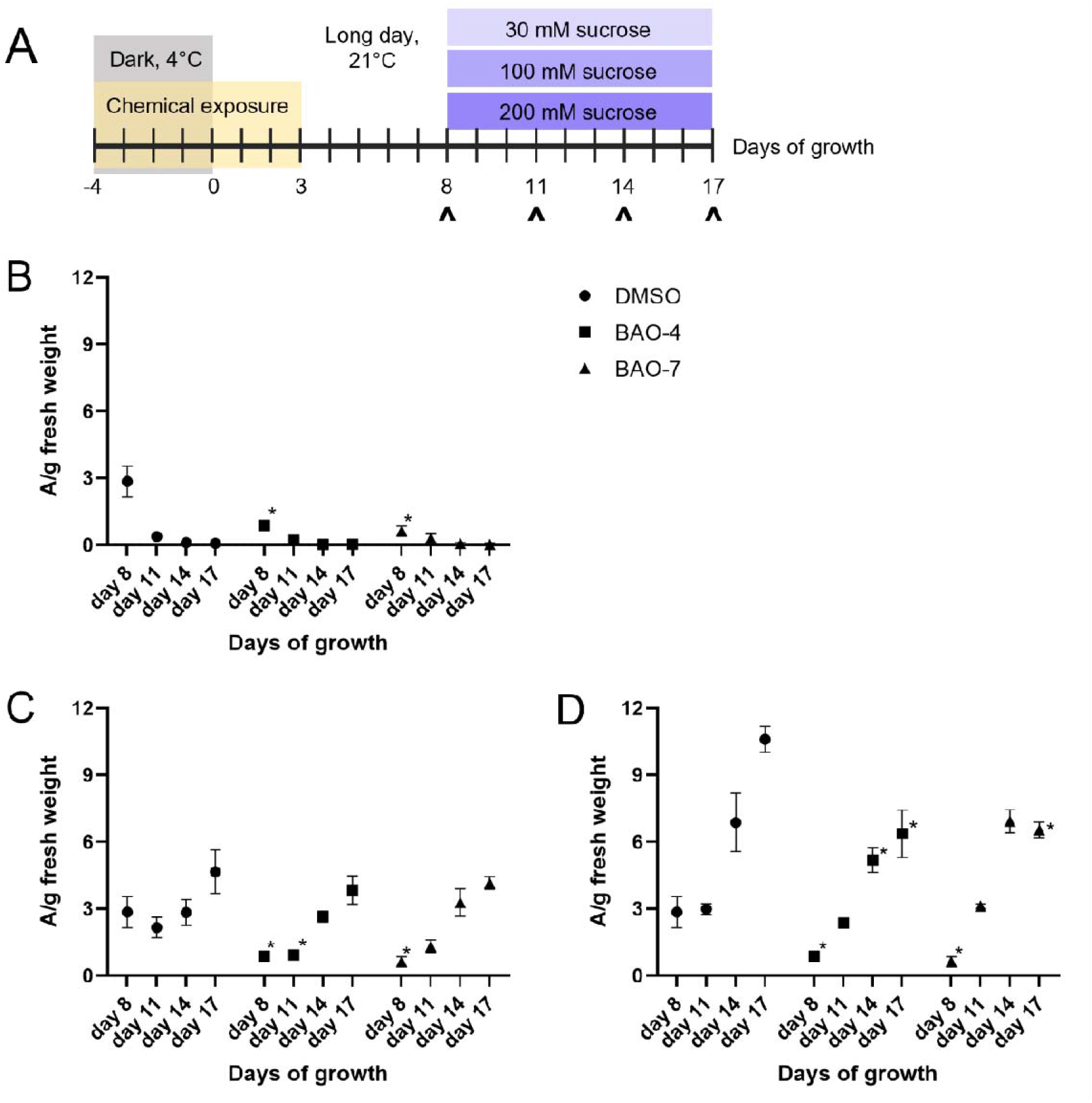
Benzaldehyde oxime chemical priming treatments can attenuate anthocyanin accumulation induced by high levels of exogenous sucrose, with temporal dynamics dependent on sucrose dose. (A) Experimental timeline indicating timeline for chemical priming exposure, recovery, sucrose application to Col-0 wild-type seeds with observation time points indicated(∧). Anthocyanin content per fresh weight observed over 12 days for seedlings chemically primed with DMSO (control), BAO-4 and BAO-7 and subsequently treated from day 8 with continued 30 mM sucrose (B), 100 mM sucrose (C), and 200 mM sucrose (D). Error bars ± SD, n = 3 pooled samples of 20-30 seedlings each. Significance of differences between BAO treatment and DMSO control treatment at each timepoint assessed with two way ANOVA, p<0.05 (*) indicated.

## Acknowledgments

CAGEF at the University of Toronto: for design and provision of the chemical genomic library. Eshan Naik: for development of the low nitrogen microtiter plate assay. Dr. Trisha Mahtani: for data collection of morphometric data replicates. Dr. Michael Stokes: for contribution to initial conceptualization and guidance.

## Supporting information

**S1 Fig. Visible anthocyanin development in liquid MS-grown *A. thaliana* seedlings supplemented with sucrose and exposed to low temperature, used for phenotypic screening**.

**S2 Fig. Example of qualitative high-throughput hit identification for chemical seed priming candidates**.

**S3 Fig. Effect of chemical priming treatment on additional growth and reproductive parameters (A) Inflorescence length (B) S1 germination percentage**.

## Notes

### Competing Interest Statement

The authors have declared no competing interest.

