## Supplementary figures and images for "A chemical genetic screen uncovers novel seed priming agents capable of persistent perturbation of anthocyanin regulation in *Arabidopsis thaliana*"

### Supplemental Figure 1

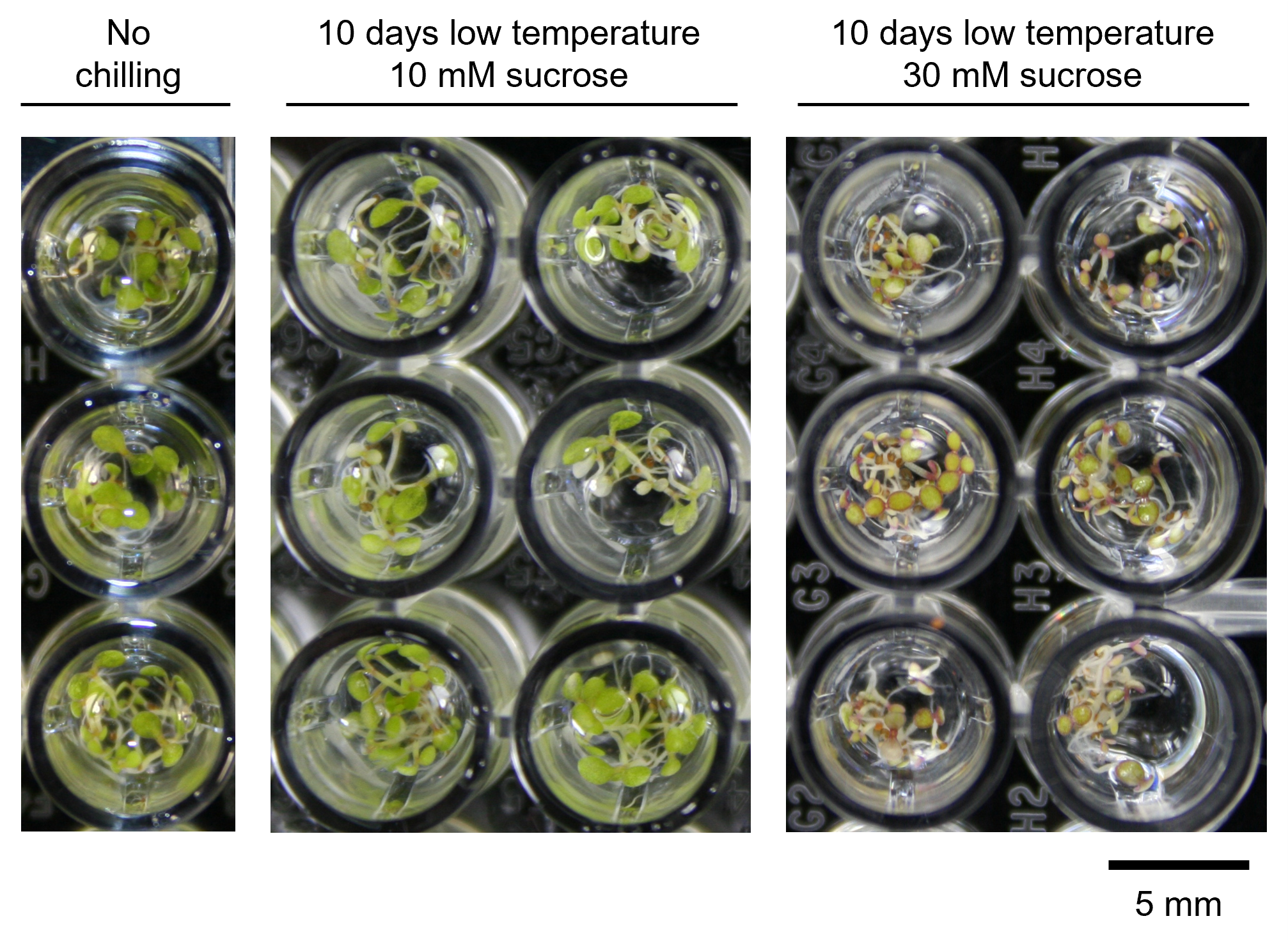

### Supplemental Figure 2

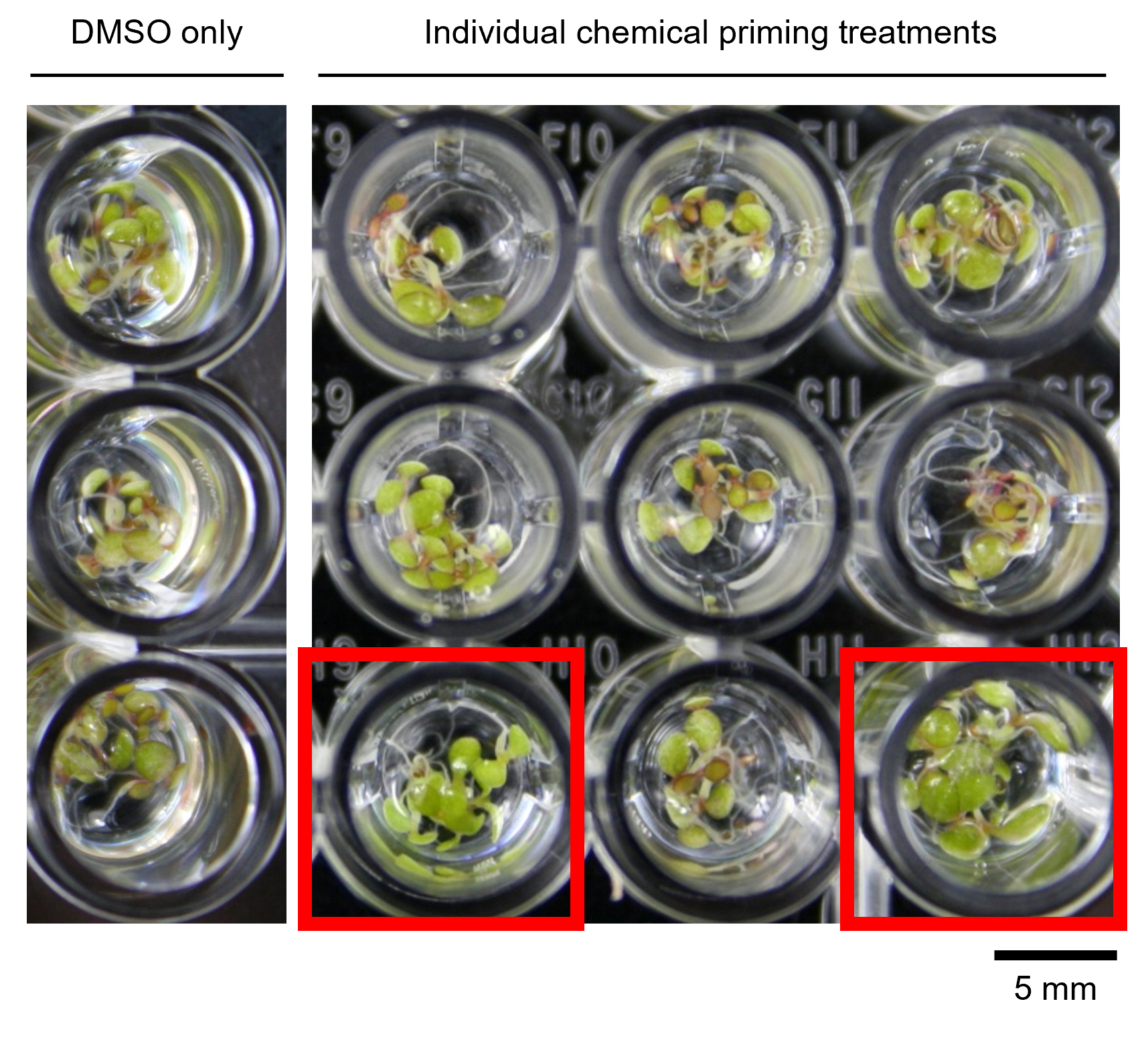

### Supplemental Figure 3

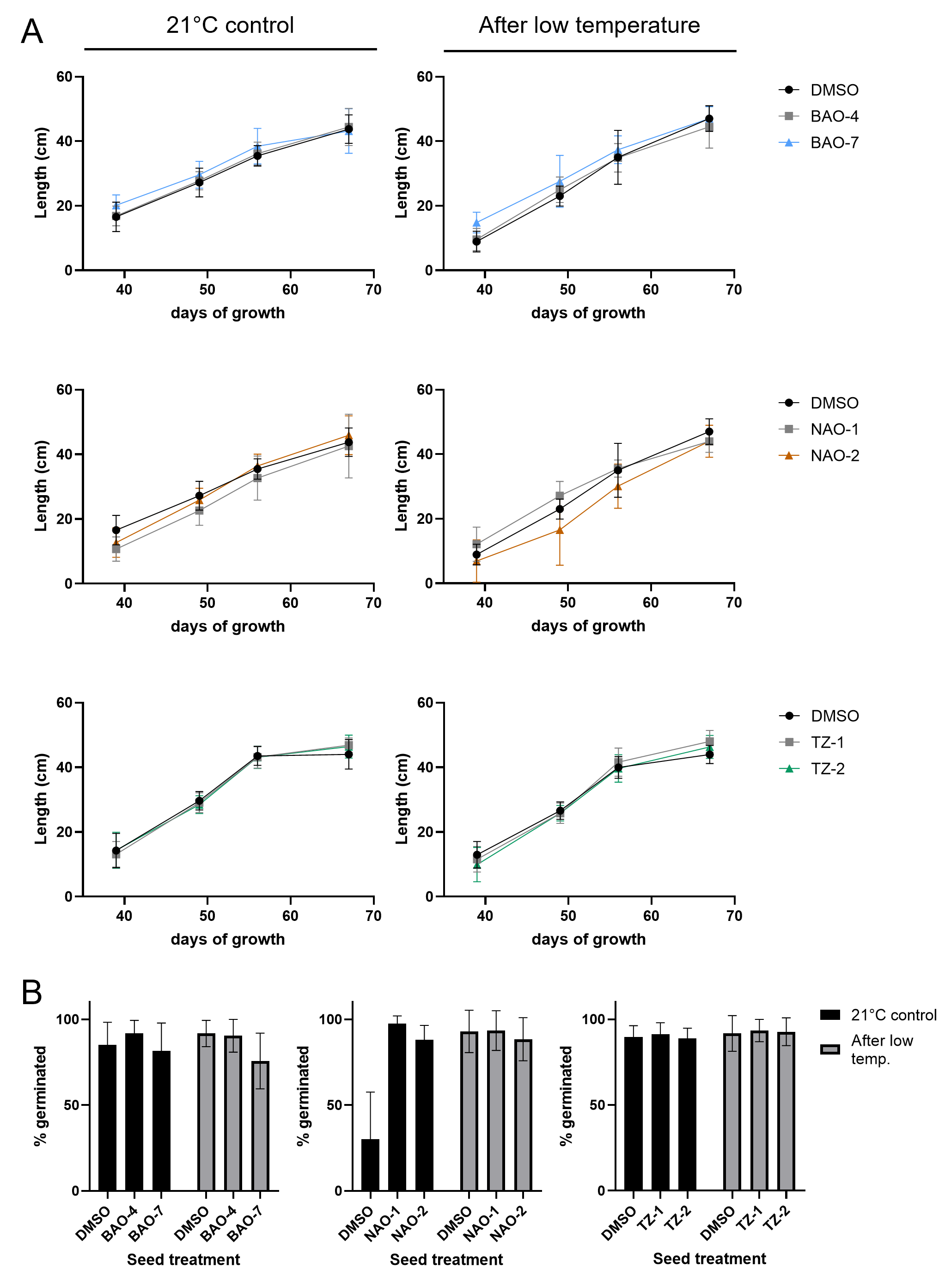
